# Model Validation Protocols for Machine Learning in Small Molecule Drug Discovery

**DOI:** 10.64898/2026.08.19.745868

**Authors:** Srijit Seal, Akshat Shirish Zalte, David Alencar Araripe, Renan Augusto Gomes, Deepa Korani, Mrinal Shekhar, Vishal Babu Siramshetty, Arijit Patra, Zhongyu Mou, Xiang Yu, Daniel Kuhn, Nils Weskamp, Jeremy Ash, Alan C. Cheng, Cheng Fang, Daniel Price, Matteo Aldeghi, Raquel Rodríguez-Pérez, Djork-Arné Clevert, Ola Engkvist, Kristine Deibler, David Rouquie, Michael Reutlinger, Nicola J. Richmond, Jon Ainsley, Mark Ledeboer, William H. Green, Andreas Bender, Cas Wognum

## Abstract

Machine learning (ML) models for molecular property prediction are increasingly deployed in drug discovery, yet their adoption in real-world scenarios requires an understanding of the conditions in which a model succeeds or fails. While standardized benchmarks are powerful instruments to measure and unlock progress in ML research, they should not be blindly treated as the end goal. Especially static and retrospective benchmarks, in which no true unknown test set is employed, limit our ability to robustly validate a model’s performance. Building on the collective expertise of a cross-industry consortium, we present a model validation framework consisting of five recommendations that would enable the community to move beyond aggregate metrics toward understanding where and why molecular property prediction models fail. We connect evaluation choices to real-world applications and case studies encountered in pharmaceutical research. The framework proposes splitting strategies that mimic realistic distribution shifts and expose common failure modes. We apply the recommended framework on a recently released dataset of absorption, distribution, metabolism, and excretion (ADME) properties. Across two complementary model algorithms, our case studies reveal four distinct failure modes (extrapolation, interpolation, representation, and evaluation) showing that model errors arise not only from distribution shift but also from limitations in molecular representations. Our results show that commonly used evaluation protocols can significantly overestimate performance and may not detect important model failure modes. All software and data are released via https://github.com/srijitseal/polaris.

## Introduction

Machine learning (ML) driven molecular property prediction is being increasingly used in early-stage drug discovery, with models influencing compound prioritization and design decisions.^1–4^ The deployment in drug discovery campaigns is a testament to the progress that has been made, yet a gap remains between perceived progress, as measured on public benchmarks, and the impact and adoption in real-world pharmaceutical applications.^5^ We argue that this gap can be partially attributed to the complexity of model validation: the structural analysis of a model’s strengths and weaknesses is application-dependent and requires interdisciplinary expertise. Unlike other ML domains, like Computer Vision or Natural Language Processing, failure modes are unintuitive and require scientific expertise or experimental validation to detect. This complexity, combined with incentive structures that reward, or even require, state-of-the-art results on existing benchmarks,^6^ has led to an overreliance on static and retrospective benchmarks. To bridge this gap, we propose a holistic evaluation framework that equips ML researchers with actionable model validation guidelines designed to mimic real-world drug discovery scenarios. This proposed framework establishes a foundation for developing robust methods and accelerating their translation into industrial pipelines.

Relying on a static and retrospective benchmark is principally problematic because it restricts the statistical replicability of experimental findings. Prior work by Ash et al. addressed replicability through repeated cross-validation and statistical tests, but public benchmarks typically suffer from another critical limitation: a lack of generalizability.^7^ ML models are prone to exploit spurious correlations that lose their predictive power in an out-of-distribution deployment, that is, when data comes from a different distribution than the training set. This is especially pronounced in drug discovery, where existing, public datasets are heterogeneous with non-obvious biases^8^ and deployment is explicitly designed to go out-of-distribution. As drug discovery campaigns progress, newly generated compounds typically explore regions of chemical space increasingly distant from the training data, while property optimization can actively drive the target distribution beyond the bounds of the original training set. During deployment, prediction errors are observed even in well-performing models, which highlights the need for understanding model failure modes. Uncertainty quantification can also help to identify when a prediction is trustworthy, and in recent work, Parrondo-Pizarro et al. analyzed uncertainty quantification for molecular ML property predictions under various dataset shifts in terms of the chemical space and property values, and activity cliffs.^9^

Due to the growing mismatch between the training distribution and the compounds encountered during deployment, predictive performance systematically degrades. This challenge motivates the framework, which is organized around four recurring failure modes. Two stem directly from the distribution mismatch. First, extrapolation failures, where models struggle when compounds are structurally distant from the training set or belong to entirely new chemical series, resulting in poor model generalization beyond its applicability domain.^10,11^ Second, interpolation failures, where models fail at activity cliffs, where minor structural changes cause massive shifts in biological activity.^12–15^ The other two stem from how molecules are encoded and how models are evaluated. Third, representation failures where fixed molecular representations are blind to activity-relevant differences such as stereochemistry, while over-predicting the effect of resonance-form changes, which correspond to the same molecule, and minor substituent changes, which leave activity largely but not entirely unchanged..^16^ Fourth, evaluation failures where validation strategies that do not match the deployment scenario, such as random or naive scaffold splits, yield misleadingly optimistic performance estimates.^17,18^

In ML, dividing a dataset into distinct training, validation, and test splits is the standard approach to evaluate how well a model generalizes to unseen data. While the field has largely adopted scaffold-based splits over random splits as a better proxy for generalization, scaffold splits are not automatically deployment-relevant^18,19^; their usefulness depends on the scaffold definition, the dataset’s scaffold-size distribution, and the extent to which the structural separation induced by the split mimics the targeted deployment scenario.^20^ For example, Fooladi et. al. benchmarked 14 classical ML and modern graph neural network (GNN) models across eight molecular datasets using ten strategies for creating out-of-distribution test sets, and found that scaffold splitting produced relatively mild distribution shifts and often overestimated generalization, whereas Unified Manifold Approximation and Projection (UMAP) clustering and Lo-Hi splits^21^ were substantially more challenging^22^; GNNs were generally somewhat more robust than classical models. Performance on standard in-distribution (ID) tests predicted out-of-distribution (OOD) performance only for easy splits, so evaluation and model selection, they conclude, should use application-relevant chemical-space splits and multiple metrics. The “right” validation strategy is context-dependent, and the field lacks a practical consensus on how to connect splitting strategy choices to specific deployment scenarios.

This context-dependence surfaces a fundamental challenge: no single benchmark can be universally applicable. ML models are deployed in different organizations that use different operational models. While standardized benchmarks remain a powerful means to track and unlock progress in ML research, they must not be blindly treated as the end goal. To improve translational use, researchers must move beyond aggregate benchmark scores and actively probe a model’s failure modes.

In this work, we build on the collective expertise and experience of a cross-industry consortium to propose a framework for model validation in molecular property prediction. The framework is summarized as five practical recommendations in Box 1, mapped to common drug-discovery deployment scenarios in Table 1, and schematized in Figure 1. Together, these elements organize model validation around a central principle: evaluation should start from the intended deployment scenario, use splitting strategies that reproduce the relevant distribution shift, verify that the intended shift was actually created, and then probe the failure modes most likely to affect real-world use. We demonstrate this framework on the Expansion Therapeutics absorption, distribution, metabolism, excretion, and toxicity (ADMET) dataset,^23,24^ a public real-world dataset combining temporal ordering (ordinal compound identifiers), chemical series structure (concentrated Butina clusters), multi-endpoint coverage (9 ADME endpoints), and multi-CRO provenance from actual drug discovery campaigns targeting RNA with small molecules. We evaluate two complementary modeling approaches: a classical XGBoost model with engineered molecular features and a graph neural network-based molecular foundation model.^25^ The framework is followed step by step by: first characterizing the dataset, then constructing and diagnosing deployment-relevant splits, and finally using those splits to expose extrapolation, interpolation, representation, and evaluation failures. Results show that commonly used evaluation practices can significantly overestimate performance, without detecting relevant failure modes. In conjunction with our open-source code, we hope that this framework will establish the foundation for a more deployment-oriented paradigm in model validation.

**Figure 1.**
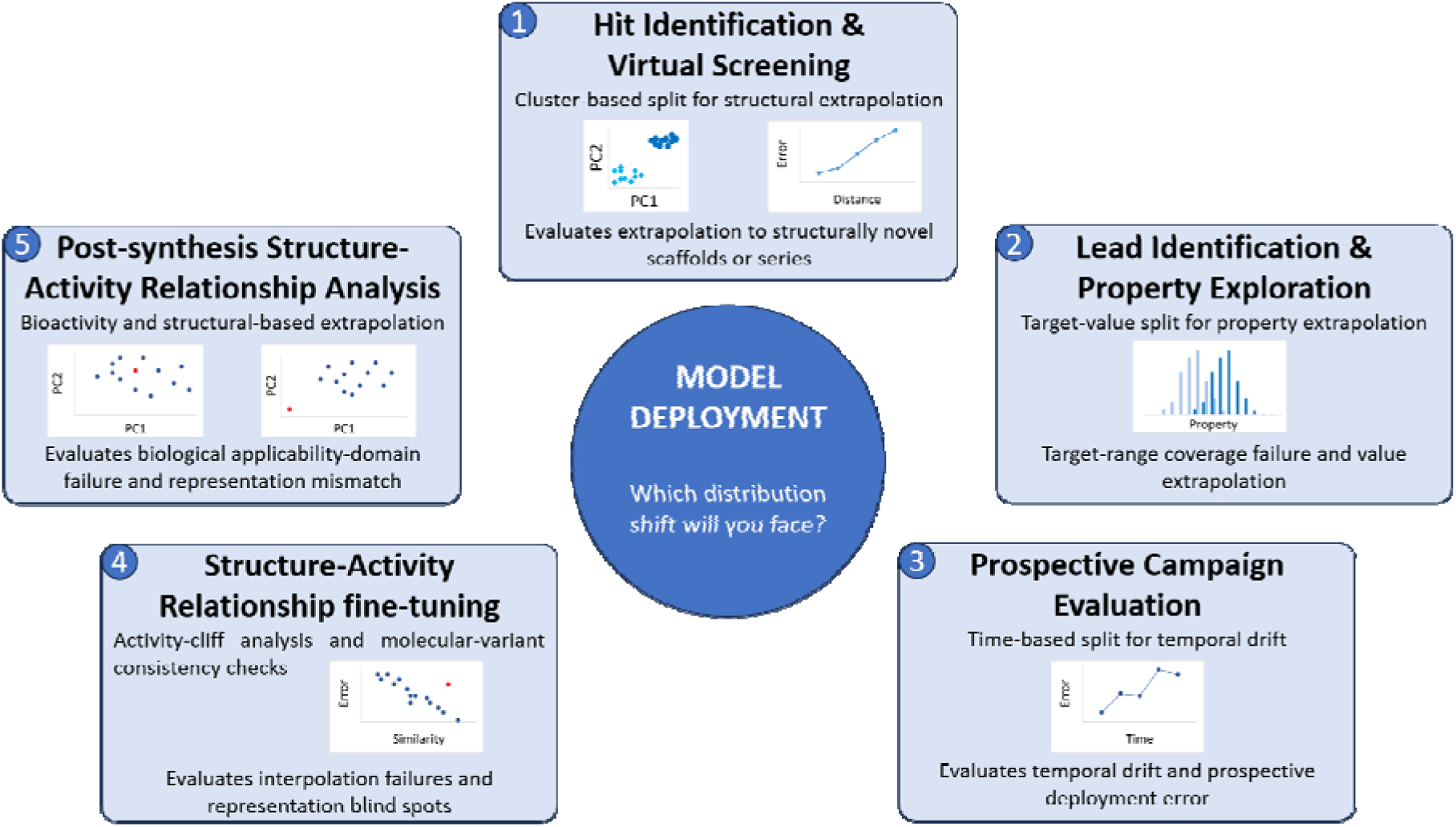
Overview of the proposed framework. Five drug discovery deployment scenarios are associated with their respective distribution shifts. The framework systematizes the selection of a suitable evaluation strategy that targets the root cause of each shift.

**Table 1.**
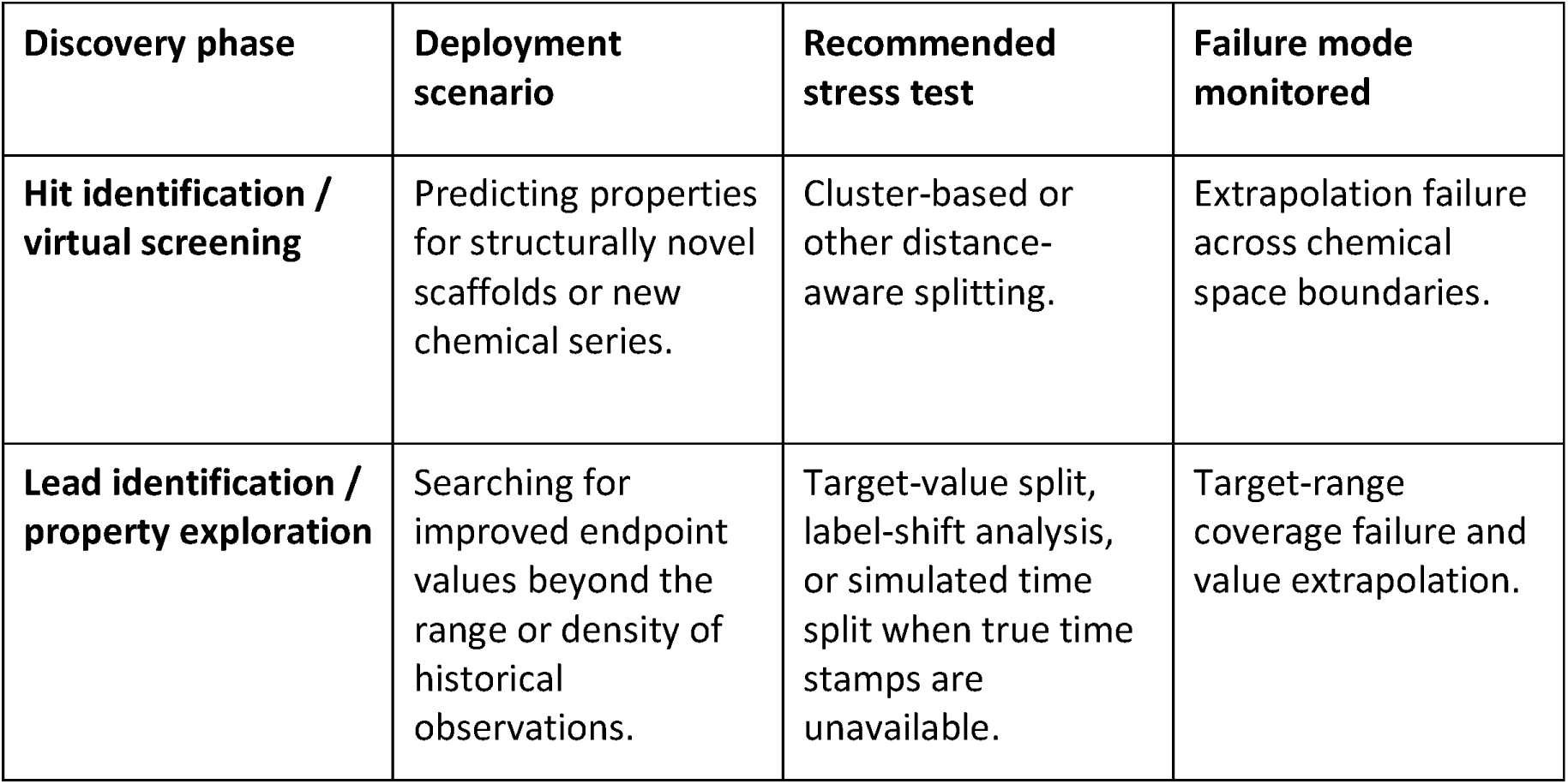
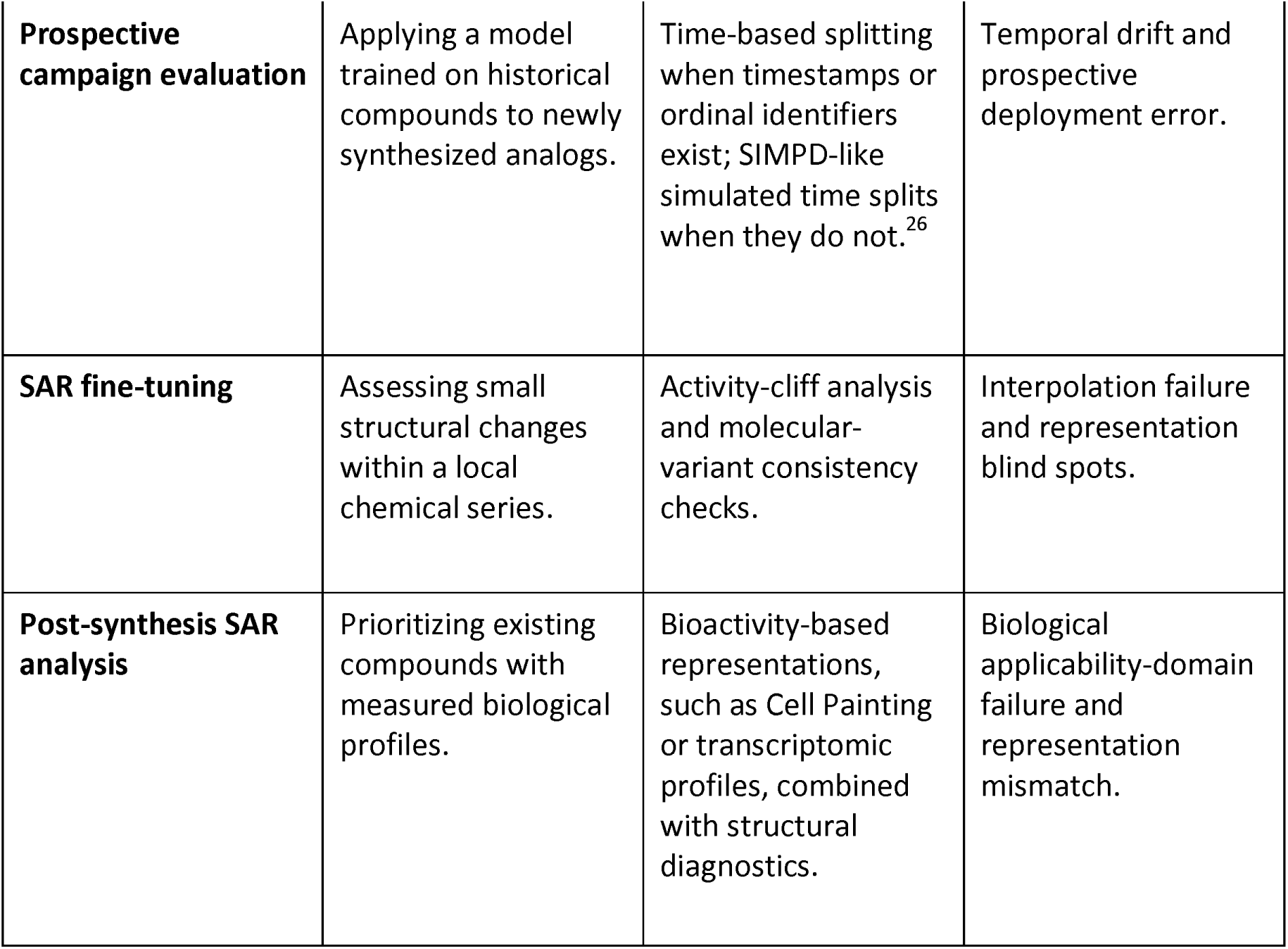
Splitting strategy and failure-mode mapping by drug-discovery phase.

| Discovery phase | Deployment scenario | Recommended stress test | Failure mode monitored |
| --- | --- | --- | --- |
| Hit identification / virtual screening | Predicting properties for structurally novel scaffolds or new chemical series. | Cluster-based or other distance-aware splitting. | Extrapolation failure across chemical space boundaries. |
| Lead identification / property exploration | Searching for improved endpoint values beyond the range or density of historical observations. | Target-value split, label-shift analysis, or simulated time split when true time stamps are unavailable. | Target-range coverage failure and value extrapolation. |
| **Prospective campaign evaluation** | Applying a model trained on historical compounds to newly synthesized analogs. | Time-based splitting when timestamps or ordinal identifiers exist; SIMPD-like simulated time splits when they do not.<sup>26</sup> | Temporal drift and prospective deployment error. |
| **SAR fine-tuning** | Assessing small structural changes within a local chemical series. | Activity-cliff analysis and molecular-variant consistency checks. | Interpolation failure and representation blind spots. |
| **Post-synthesis SAR analysis** | Prioritizing existing compounds with measured biological profiles. | Bioactivity-based representations, such as Cell Painting or transcriptomic profiles, combined with structural diagnostics. | Biological applicability-domain failure and representation mismatch. |

### Box 1.

#### Practical recommendations.

1. **Characterize the dataset before splitting.** Once the deployment scenario is specified, audit whether the available dataset contains the chemical-series structure, endpoint coverage, temporal ordering (or SIMPD-like simulated time splits26), target-value distributions, and distance distributions needed to emulate that scenario.
2. **Identify the deployment scenario.** Identify which distribution shift the model will face, and choose a splitting strategy that creates that shift by construction.
3. **Use distance-aware splits with multiple repeats.** Report confidence intervals (CIs); avoid using random splits as evidence of deployment performance when structural novelty is the use case.
4. **Verify the split actually creates the intended shift.** Check fold balance, endpoint-value distributions, nearest-neighbor distances, and structural overlap; adversarial validation can provide a complementary classifier-based diagnostic of train/test distinguishability.38
5. **Probe specific failure modes.** Performance-over-distance curves, ID/OOD series tests, activity-cliff analysis, and molecular-variant consistency each reveal blind spots that aggregate metrics hide.

## Results

In a prospective validation study, identifying the deployment scenario means defining the intended use case first and then designing the split and diagnostics around the distribution shift expected in deployment. In the present work, however, we demonstrate the framework through a case study on a fixed retrospective public dataset and ask how models trained on historical ADME measurements from medicinal chemistry campaigns would perform when used to prioritize future compounds, including new or weakly represented chemical series, molecules synthesized later in a campaign, and compounds intended to improve endpoint values beyond the historical range.

Under this deployment framing, dataset characterization is the starting point of validation as a feasibility check. The dataset must contain the structural, temporal, endpoint, and target-value structure needed to emulate the relevant deployment shifts. Accordingly, we analyse the Expansion Therapeutics ADMET dataset under the scenarios in **Table 1** (chemical-series extrapolation, prospective temporal drift, target-value extrapolation, local structure-activity relationship (SAR) discontinuities, and molecular-representation sensitivity). Further, we construct deployment-relevant splits, verify that those splits create the intended shifts, and use targeted diagnostics to expose extrapolation, interpolation, representation, and evaluation failures.

### Dataset characterization: Assay cascades create coupled data size and diversity gaps across endpoints

In pharmaceutical ADME workflows, assays are organized in a cascade: lipophilicity and solubility are measured for nearly every compound early in a program, while specialized assays such as brain protein binding are reserved for compounds that have already passed earlier property gates or when additional liabilities have been detected. This cascade structure has two consequences for predictive modeling that amplify each other. First, datasets from later-stage endpoints have substantially fewer measurements: in the Expansion Therapeutics dataset (7,608 ML-ready molecules, 9 ADME endpoints spanning lipophilicity, solubility, metabolic stability in liver microsomes, permeability, and protein binding; Table 2), coverage ranges from 96% (LogD, KSOL) to 6% (MGMB). Second, we calculated Jaccard distances to the nearest neighbor (1-NN) using Morgan (ECFP4) fingerprints. The within-endpoint median 1-NN Jaccard distance is tight across all endpoints (0.18–0.20, Table 2), indicating that every endpoint subsample is drawn from structurally similar chemical space rather than differing in local density; the generalization challenge therefore comes from the reduction in sample size at later-stage endpoints rather than a drop in local structural coverage. The label matrix is also incomplete: each molecule has measurements for a median of 4 out of 9 endpoints. All 9 endpoints show statistically significant statistically significant train/test distribution shift (though effect sizes vary widely, D = 0.04–0.37; two-sample Kolmogorov–Smirnov (KS) test, chosen because endpoint value distributions are non-normal; Table 2), with Caco-2 Papp A>B (KS D statistic= 0.28) and Caco-2 Efflux (KS D statistic = 0.37) exhibiting the strongest shift, consistent with permeability assays entering the cascade later as compounds advanced. Endpoints that are measured late-stage consequently have fewer available measurements, although their position in the assay cascade does not necessarily indicate greater decision impact. Differences in endpoint coverage and distribution shift should therefore be interpreted in the context of each assay’s purpose within the experimental cascade. Accurately predicting assays that are decision-critical could front-load risk and lower failure rates and thus costs.

**Table 2.**
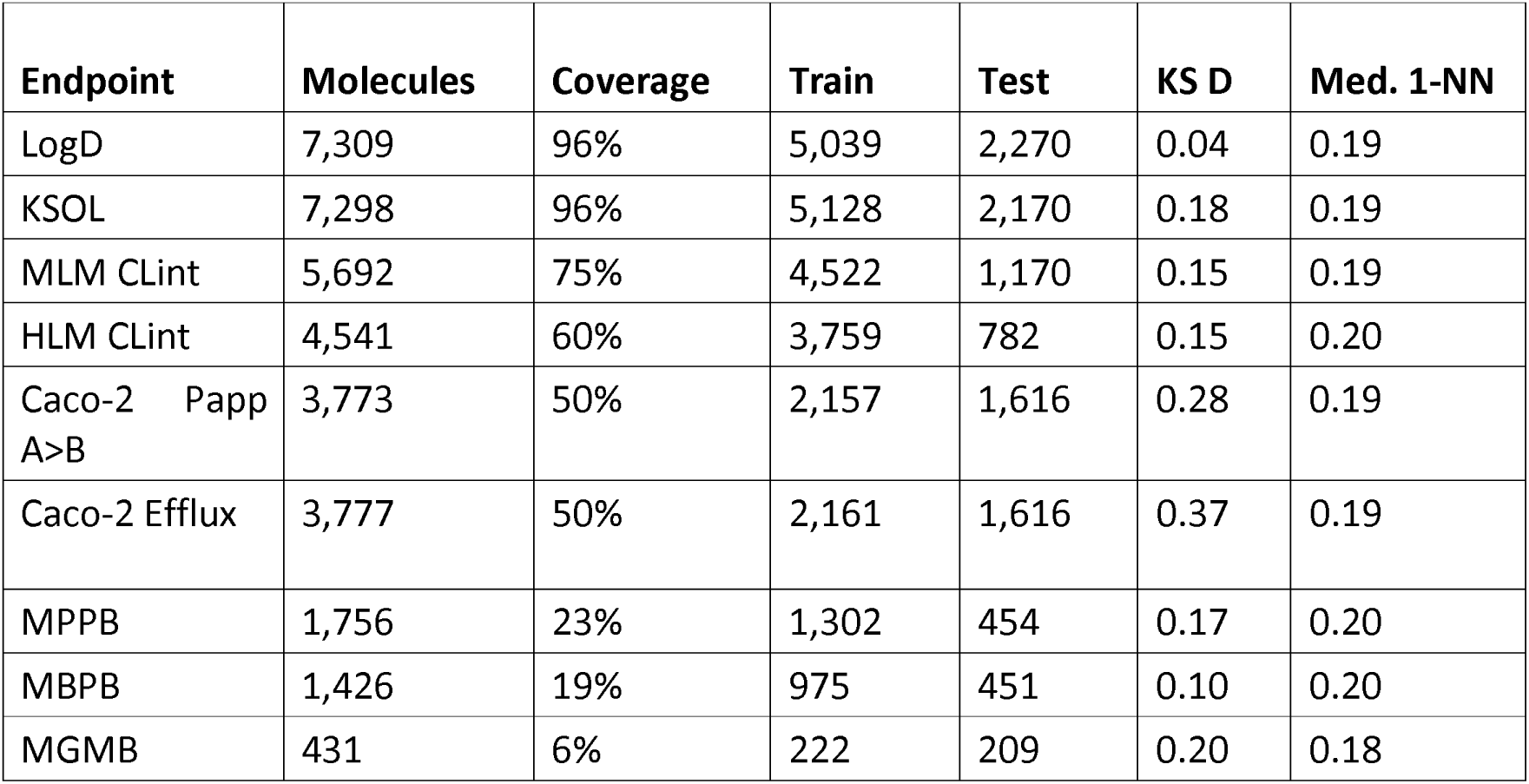
Endpoint summary for the Expansion Tx dataset. Coverage: fraction of the 7,608 ML-ready molecules with a non-null measurement for that endpoint. The Kolmogorov-Smirnov D statistics (KS D) comparing train and test endpoint value distributions and Med. (1-NN: median nearest-neighbor Jaccard distance within each endpoint’s measured compounds (lower values indicate more structurally concentrated chemistry) are shown.

| Endpoint | Molecules | Coverage | Train | Test | KS D | Med. 1-NN |
| --- | --- | --- | --- | --- | --- | --- |
| LogD | 7,309 | 96% | 5,039 | 2,270 | 0.04 | 0.19 |
| KSOL | 7,298 | 96% | 5,128 | 2,170 | 0.18 | 0.19 |
| MLM CLint | 5,692 | 75% | 4,522 | 1,170 | 0.15 | 0.19 |
| HLM CLint | 4,541 | 60% | 3,759 | 782 | 0.15 | 0.20 |
| Caco-2 Papp A>B | 3,773 | 50% | 2,157 | 1,616 | 0.28 | 0.19 |
| Caco-2 Efflux | 3,777 | 50% | 2,161 | 1,616 | 0.37 | 0.19 |
| MPPB | 1,756 | 23% | 1,302 | 454 | 0.17 | 0.20 |
| MBPB | 1,426 | 19% | 975 | 451 | 0.10 | 0.20 |
| MGMB | 431 | 6% | 222 | 209 | 0.20 | 0.18 |

A physicochemical property analysis shows that the median molecular weight (MW) is 383 Da (mean 393 Da; only 3.5% exceed 500 Da), median MolLogP is 3.33, median HBA is 4 (only 0.6% exceed 10), median HBD is 1, median FractionCSP3 is 0.36, and the median molecule carries 3 aromatic rings. Combining the four core Lipinski rules (MW ≤ 500, LogP ≤ 5, HBA ≤ 10, HBD ≤ 5), 93.5% of the dataset passes all four, showing the dataset largely comprises drug-like molecules, as expected.

To contextualize the structural novelty of the Expansion Therapeutics compounds, we computed the maximum Tanimoto similarity of each dataset molecule to ChEMBL 36 (2.85M compounds) and its ADME-annotated subset (308K compounds). The median maximum similarity was 0.44 to all of ChEMBL and 0.39 to the ADME subset, with 25% of compounds having no ChEMBL neighbor above 0.4 similarity and 98% below the 0.7 activity-relevant threshold (Figure S1). This confirms that the dataset occupies relatively underexplored chemical space.

The OpenADMET challenge competition used a time-split, where we found test molecules to be systematically farther from the training set than the training molecules were from each other. Specifically, the test-to-train 1-NN median distance was 0.37, nearly double the within-train median of 0.20 (Figure 2). This gap reflects the presence of compounds that lie outside the chemical space covered by the training set, rather than a strictly monotonic change over time. In medicinal chemistry workflows, early-stage exploration typically involves diverse scaffolds, while later-stage optimization focuses on closely related analogues within established chemical series. While it may seem random split would work later-stage optimization because projects are looking at analogs, even on a per project basis, it can be overoptimistic compared to temporal splits per-project.^27^ Overall, across an entire campaign, the introduction of new series and continued exploration results in a dataset that combines both local refinement and broader expansion of chemical space.

**Figure 2.**
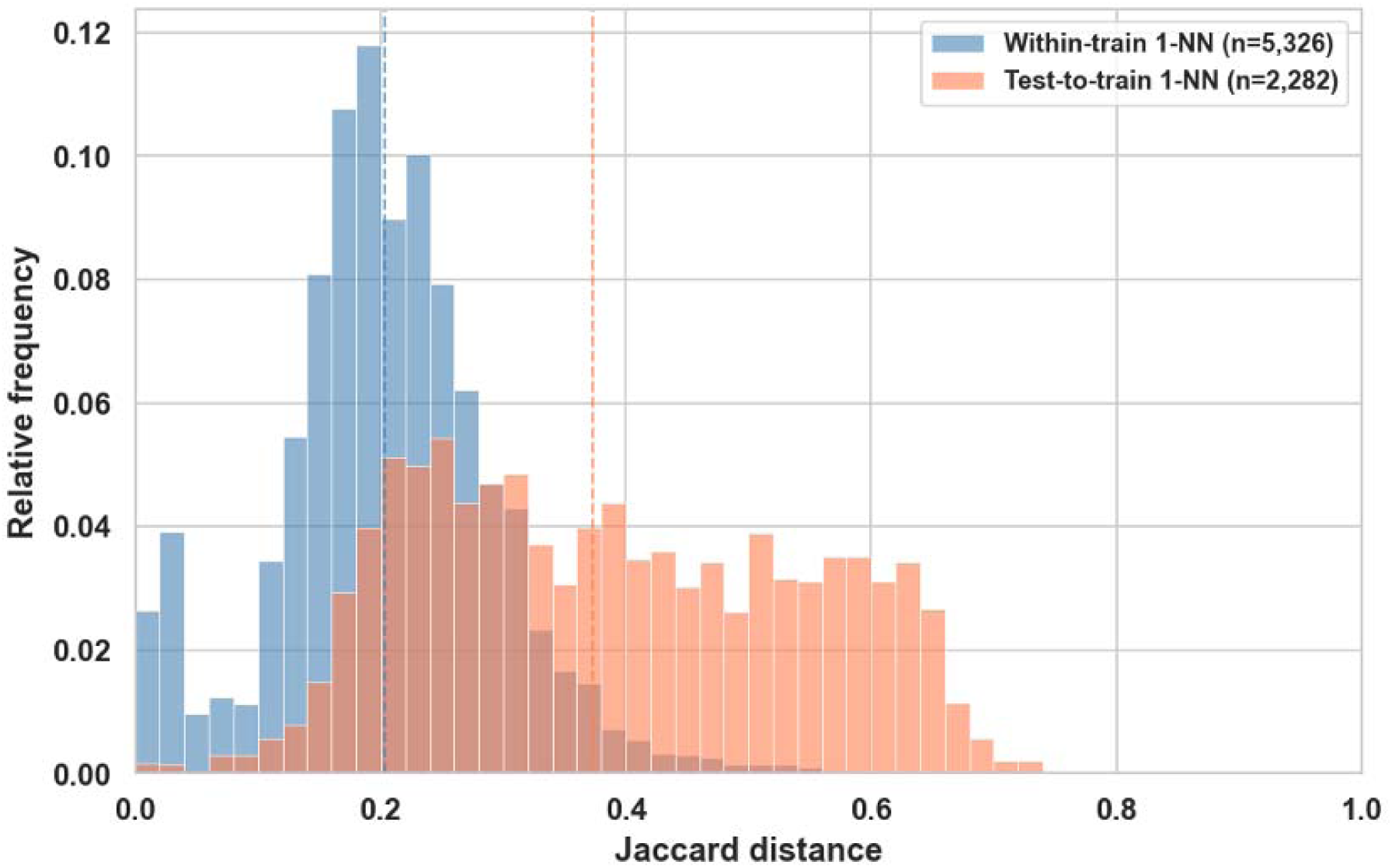
Test-to-train versus within-train 1-NN Morgan fingerprints Jaccard distance distributions, showing the structural gap between training and test sets in the competition split.

Further, in drug discovery programs, the majority of medicinal chemistry effort is concentrated on a few promising chemical series, while smaller exploratory series and singleton screening hits populate the long tail. Many series and singletons of hits are not further explored; instead, effort tends to concentrate on a few closely related congeneric series comprising thousands of compounds. We characterized this setup in the Expansion Therapeutics dataset using Butina clustering at a Jaccard distance cutoff of 0.7, which identified 135 clusters. Three dominant clusters containing 2,572, 1,301, and 885 molecules together comprise 62.5% of the entire dataset (Figure 3), with the remaining molecules spread across 132 smaller clusters, with a median size of 4, including 33 singletons (24.4%). A model fit to this distribution therefore learns predominantly from a handful of series and must extrapolate to the long tail, the regime it operates in once deployed against compounds not yet synthesized.

**Figure 3.**
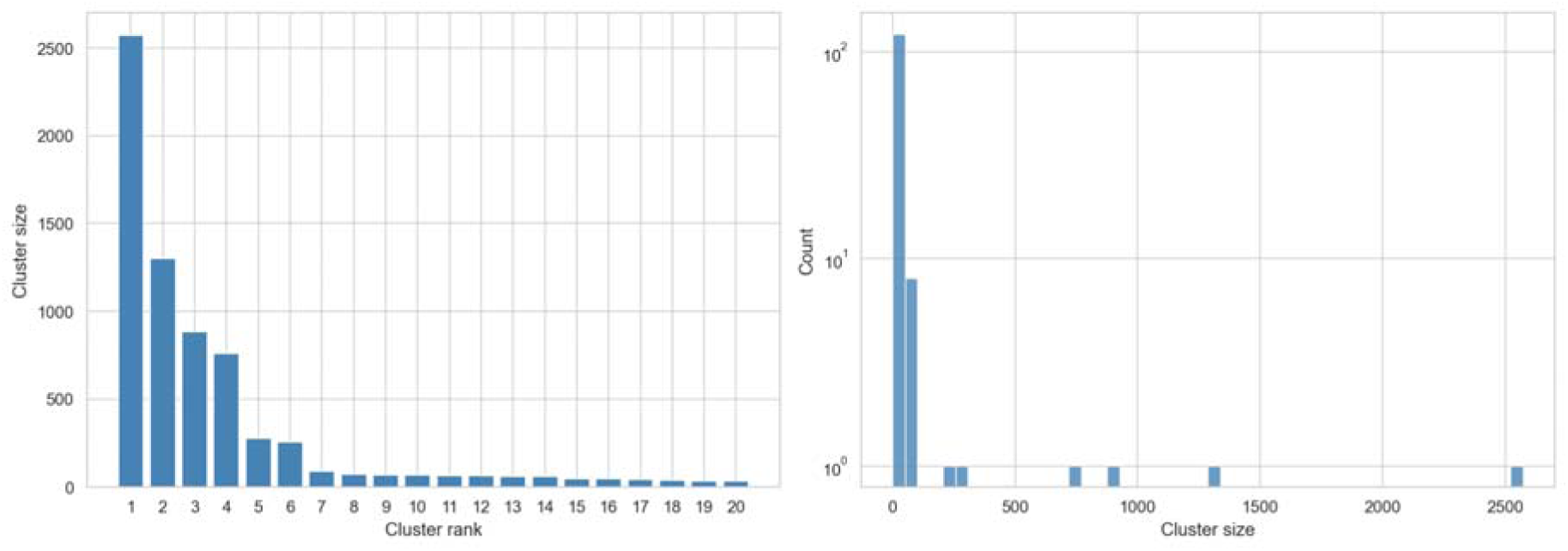
Butina cluster size distribution (Jaccard distance cutoff 0.7). Left: cluster size by rank for the 20 largest clusters. Right: histogram of all 135 cluster sizes (log scale); 33 are singletons. Three dominant clusters contain 62.5% of all molecules, reflecting the concentrated chemical series structure of the drug discovery programs in the Expansion Therapeutics dataset.

The characterisation here is specific to the dataset. Applying the same characterization to the public Biogen ADME dataset^28^(which is more global in nature across various programs) as shown in Figure S2, shows it lacks concentrated chemical-series structure: its largest Butina cluster spans only 2.1% of compounds (versus 33.8% here) and its top ten clusters cover just 14% (versus 84%), with 54% of clusters being singletons. As a consequence a series-based ID-vs-OOD split is not constructible as such, and even a random split already separates test from training as strongly as a deliberate split does in ExpansionRx (median test-to-train 1-NN distance 0.57 vs 0.20). This underscores that the appropriate split depends on the dataset and is established by characterization, rather than a protocol applied blindly. In pharmaceutical settings, the deployed ADME models learn from data across several programs, and these are usually denominated “global” models as they learn from a broad chemical space and historical data. Sometimes global ML models are also fine-tuned on specific project data.^27,29^

### Splitting strategies probe complementary generalization axes

The choice of splitting strategy should be driven by the deployment scenario. In hit identification campaigns, where compound libraries are screened for activity against a new target, the model encounters scaffolds absent from its training data, making cluster-based evaluation essential. In lead optimization, medicinal chemists iteratively modify a promising series to improve multiple specific properties. In this context, the relevant question is whether the model can extrapolate beyond the historical value range, making target-value splits appropriate. In prospective deployment, where a model trained on a program’s historical compounds is applied to newly synthesized analogs, time-based splits most faithfully represent the temporal distribution shift. No single split captures all deployment realities, and the models should be evaluated in multiple complementary splitting methodologies.

In this work, we studied cluster-based, time-based, and target-value splitting for cross-validation. The three splitting strategies produced distinct test-to-train distance distributions, illustrated here for LogD as a representative endpoint (Figure 4), confirming they test different generalization challenges. Cluster-based splitting, as expected, produces large structural distances (median 1-NN 0.37–0.44 across 5 folds), mimicking the scenario of predicting properties for a novel chemical series not represented in the training data. Time-based splitting produced intermediate distances (0.30–0.39) with non-monotonic per-fold patterns, suggesting non-linear chemical series exploration, a pattern consistent with medicinal chemistry programs that revisit earlier scaffolds as new biological data emerges. Target-value splitting produced the lowest structural distances (0.24–0.29), testing whether models can extrapolate to value ranges not seen during training, a challenge that arises when optimizing a property beyond the range of historical compounds.

**Figure 4.**
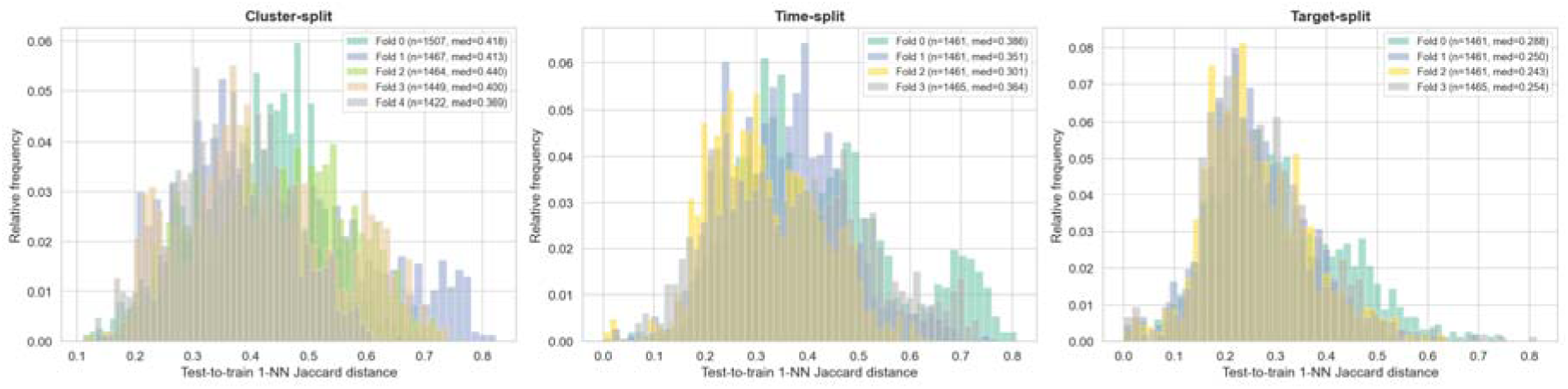
Test-to-train 1-NN Jaccard distance distributions per fold for three splitting strategies (LogD, representative endpoint). Cluster-based splitting (5 folds) produces the largest structural separation (per-fold medians 0.369–0.440), time-based splitting (4 folds, 0.301– 0.386) is intermediate, and target-value splitting (4 folds, 0.243–0.288) keeps test molecules structurally closest to training. Per-fold medians and test-set sizes are given in the legends.

Consistent with prior studies, there was also a significant label shift over time-based splits in 8 of 9 ADME assays, strongest for the Caco-2 permeability assays, kinetic solubility, and mouse liver microsomal clearance, with only mouse blood protein binding (MBPB) showing no reliable trend (Figure S3). Drift direction is assay-specific rather than uniformly optimizing^30^: LogD and clearance trend downward over time while solubility and efflux trend upward. The time-based split shifts test values within the range already seen in training (≤0.9% of test molecules fall beyond the training range in every assay), whereas genuine value extrapolation is produced solely by the target-value split (91.5% of LogD test molecules beyond range; Figure S3). Thus, temporal ordering captures drift in property space, as expected when medicinal chemistry programs iteratively optimize the properties being predicted. Overall, temporal deployment and value extrapolation are distinct generalization axes that should be reported separately rather than conflated.

To quantify the change in the molecular feature distribution between training and test sets, directly rather than through a 1-NN Jaccard distance metric, adversarial validation was implemented using a classifier trained on ECFP4 fingerprints to distinguish test molecules from their training set. Area under the Receiver Operating Characteristic Curve (AUROC) were calculated for all three splitting strategies; cluster (AUROC = 1.00), time (AUROC = 0.98), and target-value (AUROC = 0.86) splits, the test molecules typically occupy a different region of chemical space than the training molecules relative to a random split (AUROC = 0.51), indicating a covariate shift. Along with the label-shift (KS D statistic of the target distribution: cluster 0.06–0.17, time 0.06–0.27, target 0.96–0.99 across all assays), the two metrics show that cluster and time splits lead to new chemical space being evaluated (high AUROC, low KS D statistic), and the target-value split, driven by label shift, leads to lower structural separability.

These three data splits are complementary. Cluster-based splitting tests extrapolation in chemical space, time-based splitting tests robustness to temporal drift in design strategy, and target-value splitting tests the limits of value extrapolation. Together, they provide a more complete picture of generalization than any single strategy alone. All strategies produced balanced fold sizes (max/min ratio < 1.15), ensuring that performance differences reflect genuine distributional challenges rather than data imbalance artifacts.

### Failure mode 1a: performance degrades with structural distance and target-value extrapolation

A central question for model deployment is whether model error increases as test molecules become more structurally distant from training data, and if so, by how much. The previous section defines the deployment stress tests, while this section quantifies how those stress tests change model error under the two architectures.

For both XGBoost and CheMeleon models, we tracked prediction error as a function of the 1-NN distance (Morgan fingerprints, useChirality=True) under all three splitting strategies. We used Relative Absolute Error (RAE) and its macro-averaged form across all nine endpoints (MA-RAE) as primary metrics for comparing performance. Both models showed systematic performance differences across splitting strategies. Under XGBoost, MA-RAE was 0.67 (cluster), 0.77 (time), and 6.12 (target) and under CheMeleon MA-RAE values were 0.63, 0.73, and 5.26, respectively (Table 3; for test sets defined by the competition-split, performance across all endpoints in both models are shown in Figure S4, Table S1).

**Table 3.** Per-endpoint RAE under three splitting strategies, mean across folds, for both XGBoost (Optuna TPE-tuned on ECFP4 + RDKit 2D descriptors) and CheMeleon (graph-foundation model fine-tuned per endpoint). Cluster splits test structural novelty, time splits test prospective drift, and target-value splits test extrapolation beyond the observed endpoint-value range.

| Endpoint | Cluster RAE (XGB) | Cluster RAE (CheM) | Time RAE (XGB) | Time RAE (CheM) | Target RAE (XGB) | Target RAE (CheM) |
| --- | --- | --- | --- | --- | --- | --- |
| LogD | 0.50 | 0.38 | 0.58 | 0.46 | 3.68 | 2.78 |
| KSOL | 0.70 | 0.60 | 0.81 | 0.70 | 18.31 | 14.00 |
| HLM CLint | 0.83 | 0.79 | 0.82 | 0.80 | 5.55 | 4.96 |
| MLM CLint | 0.74 | 0.73 | 0.88 | 0.91 | 5.13 | 5.25 |
| Caco-2 Papp A>B | 0.69 | 0.78 | 0.74 | 0.84 | 4.99 | 4.75 |
| Caco-2 Efflux | 0.62 | 0.61 | 0.75 | 0.76 | 3.57 | 3.30 |
| MPPB | 0.65 | 0.60 | 0.78 | 0.70 | 4.84 | 3.97 |
| MBPB | 0.59 | 0.50 | 0.67 | 0.56 | 4.03 | 3.46 |
| MGMB | 0.73 | 0.65 | 0.91 | 0.82 | 4.97 | 4.87 |
| **MA-RAE** | **0.67** | **0.63** | **0.77** | **0.73** | **6.12** | **5.26** |

Performance-over-distance curves revealed root-mean square error (RMSE) degradation with structural distance under cluster-based splitting for most endpoints, that is the model’s error at the most distant test molecules worse than at the closest ones (Figure 5), with LogD showing 3.6-fold degradation under XGBoost, HLM CLint 3.04-fold, MPPB 2.14-fold, and MLM CLint 2.08-fold; under CheMeleon, HLM CLint degradation increases to 4.30-fold. Spearman rank correlations (ρ) between 1-NN distance and absolute prediction error confirm a consistent monotonic relationship across all endpoints: under cluster-based splitting, ρ ranges from 0.08 (Caco-2 Papp) to 0.26 (MGMB) for XGBoost and from 0.08 (KSOL) to 0.34 (MGMB) for CheMeleon. The moderate correlation magnitudes reflect that distance is a necessary but not sufficient predictor of error; many distant molecules are still predicted accurately, while some nearby molecules (e.g., activity cliff pairs) produce large errors despite small distances.^9^ Endpoint-specific degradation patterns showed that even properties governed by global molecular features were affected. LogD, a bulk thermodynamic property governed by the balance of polar and nonpolar surface area (as well as flexibility and size of the molecule), nonetheless degrades substantially (3.60-fold with XGBoost and 3.54-fold with CheMeleon) because the Morgan fingerprints representation changes dramatically between distant scaffolds even when global polarity is conserved.For endpoints influenced by bulk physicochemical properties, fingerprint-based models may generalize more robustly across chemical series than for endpoints sensitive to specific structural motifs. Under XGBoost on ECFP4 and RDKit two-dimensional (2D) descriptors, Caco-2 endpoints were notable exceptions showing minimal degradation (Papp A>B 0.98-fold, Efflux 0.95-fold). This performance stability over structural distance may be consistent with the biophysics of passive membrane permeability; Caco-2 Papp is influenced by molecular size, lipophilicity, and hydrogen bonding capacity, global molecular properties that are captured by the RDKit 2D descriptors and less so by scaffold identity. Under the CheMeleon foundation model, Caco-2 endpoints degrade 2.91-fold (Papp) and 1.60-fold (Efflux). HLM CLint similarly degrades 4.30-fold under CheMeleon versus 3.04-fold under XGBoost.

**Figure 5.**
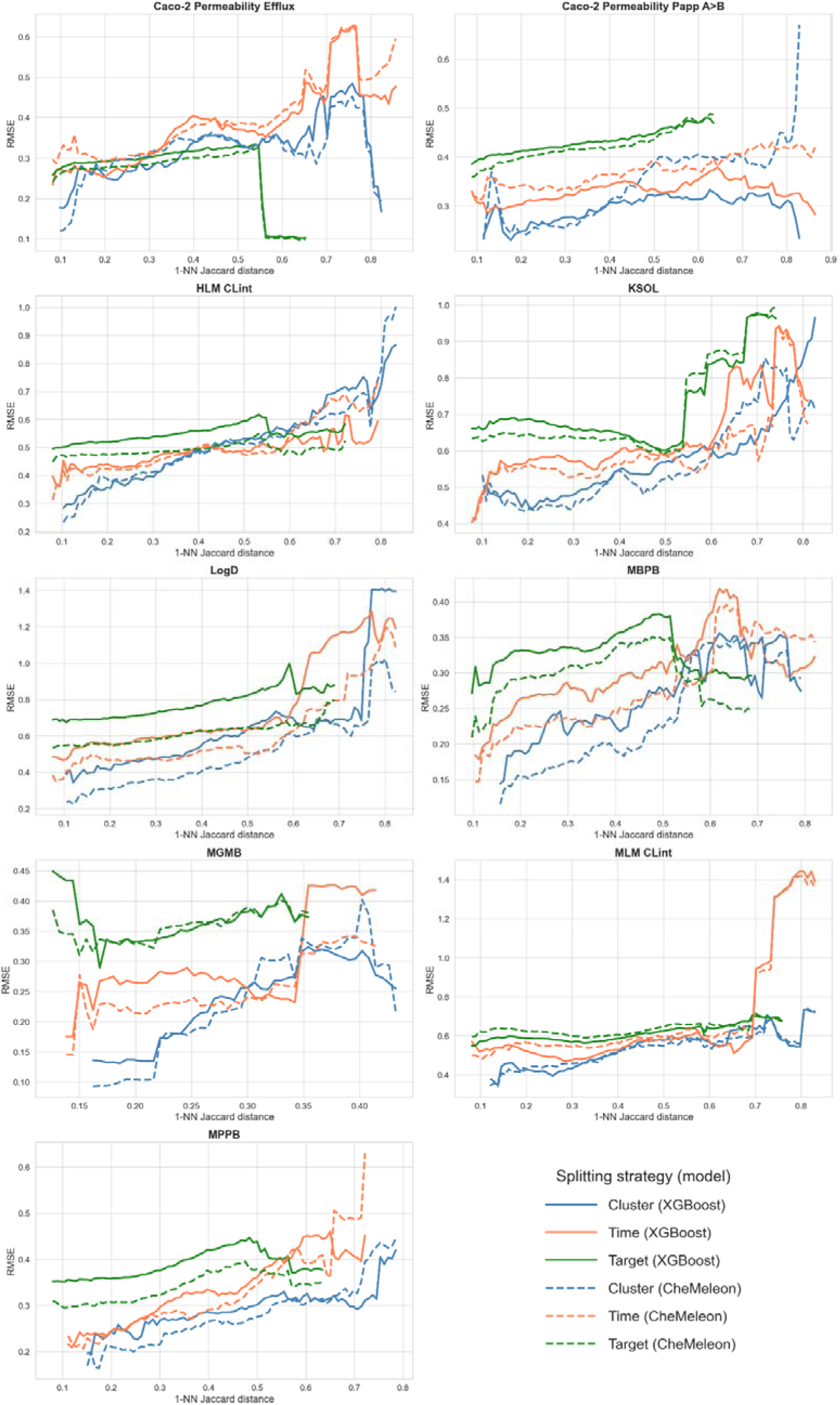
Performance-over-distance curves for all 9 endpoints across three splitting strategies, comparing XGBoost (top row of each panel) and CheMeleon (bottom row). RMSE generally increases with 1-NN Jaccard distance to training data in both models (rolling-window RMSE increases with 1-NN Jaccard distance to training data in 47 of 54 endpoint × strategy × model curves; positive Spearman rank correlation between binned distance and RMSE; significant at p < 0.05 in 49 of 54 curves).

XGBoost is a gradient boosting method whose prediction is an additive sum where the leaf weights are gradient steps but with limited extrapolation to target ranges outside training data. The extremely poor target-split MA-RAE under XGBoost (6.12) is likely because out-of-range test molecules were poorly predicted with larger errors. CheMeleon’s target-split also shows a high error of MA-RAE = 5.26. Neural network regressors are not formally bounded by the training-target range, yet without sufficient signal in the training distribution to learn how the property continues outside it, the foundation-model representation extrapolates poorly as well. Due to the lack of extreme data examples and the employed loss function, there is a general regression to mean effect in various ML models. The shared poor outcome is that target-split is by construction not a generalization test that any of the two architectures can pass in the case of this dataset; it is most useful as a diagnostic for value-range coverage rather than model-comparison.

Across all 9 endpoints, CheMeleon yields cluster-/time-/target-split MA-RAE of 0.63 / 0.73 / 5.26 (versus 0.67 / 0.77 / 6.12 for XGBoost): the strategy ranking and the qualitative degradation pattern are preserved, foundation-model representation lifts the floor on every strategy, but does not eliminate distance-driven failure.

### Failure mode 1b: ID versus OOD degradation across chemical series boundaries

We next investigated how predictive performance degrades when a model is applied to a chemical series absent from its training data, that is, cross-series generalization, as opposed to in-series generalization within a chemical series already represented in training. The two largest Butina clusters (2,572 and 1,301 molecules) were used to construct this comparison: the largest cluster was split temporally into an in-distribution (ID, in-series) training set (80%, n = 2,057) and an ID test set (20%, n = 515), while the second-largest cluster, structurally distinct and sharing no molecules with the first, served as the out-of-distribution (OOD, cross-series) test set, mirroring how historical ADME models are adapted to project-specific chemical series.28,30 Models were trained only on the ID training set and evaluated separately on the ID and OOD test sets, allowing direct comparison of within-series versus cross-series performance for the same trained model. MGMB was excluded for insufficient coverage (7–17%). By construction, ID test molecules remain structurally close to the training data (median 1-NN distance 0.29), while OOD test molecules are far more distant (0.763). We report median squared error, rather than RMSE, because it is less sensitive to the small number of very large errors expected among structurally distant OOD molecules, which would otherwise dominate an RMSE-based comparison. Notably, ID performance was not uniformly strong: HLM CLint showed negative R² even on the ID test set under both XGBoost (−0.03) and CheMeleon (−0.65), indicating that this endpoint is difficult to predict even for molecules drawn from the same chemical series as training, independent of any distribution shift.

For each endpoint, both Optuna TPE-tuned XGBoost and the fine-tuned CheMeleon foundation model were trained on the ID training set and evaluated on both the ID and OOD test sets (Figure 6, Table 4). Under XGBoost, the coefficient of determination (R²) dropped below zero for 5 of 8 endpoints on the OOD set (HLM CLint, MLM CLint, Caco-2 Papp, Caco-2 Efflux, MBPB), meaning the model explained less variance than a constant mean-predictor baseline on the held-out OOD set. The remaining endpoints retained weakly positive R² on OOD data but still degraded substantially. Median squared error increased up to 6.6-fold, with KSOL (6.6-fold), MLM CLint (6.4-fold), MPPB (5.5-fold), HLM CLint (5.2-fold), LogD (5.1-fold), and MBPB (5.1-fold). MLM CLint R² drops from 0.107 (ID) to −0.82 (OOD). By contrast, Caco-2 Papp A>B shows the smallest XGBoost median-SE degradation (1.0-fold). However, its OOD R² is still negative (−0.04), indicating that the residual error distribution changes even when the median squared error does not. Remarkably, LogD, despite being the best-performing endpoint under ID conditions (R² = 0.645), still degrades median-SE by 5.1-fold on OOD data (R² = 0.05, and the OOD set has 1.8× higher variance than the ID variance). This value means that XGBoost explains only ∼5% of the variance in the structurally distinct OOD series relative to a constant mean-predictor baseline, even though it performs well within the training series. On evaluation, we find the OOD series is chemically distinct in its basic centre: 85% of OOD compounds contain an amino-pyridine versus 97% aliphatic amines in the ID series, and the model over-predicts OOD LogD by a systematic 0.4-1.5 log units (versus a near-zero ID bias of +0.09). This directional offset is likely due to an ionization/pKa regime shift that an ID-trained model cannot capture. At the fixed assay pH (∼7.4), the fraction of each molecule that is ionized depends on its pKaH, so a model anchored on fully protonated aliphatic amines (pKaH ≈ 9-10) systematically misjudges a series built on amino-pyridines (pKaH ≈ 5-9), which are only partially ionized at that pH. The result underscores that within-series performance is a fundamentally unreliable predictor of cross-series generalization.

**Figure 6.**
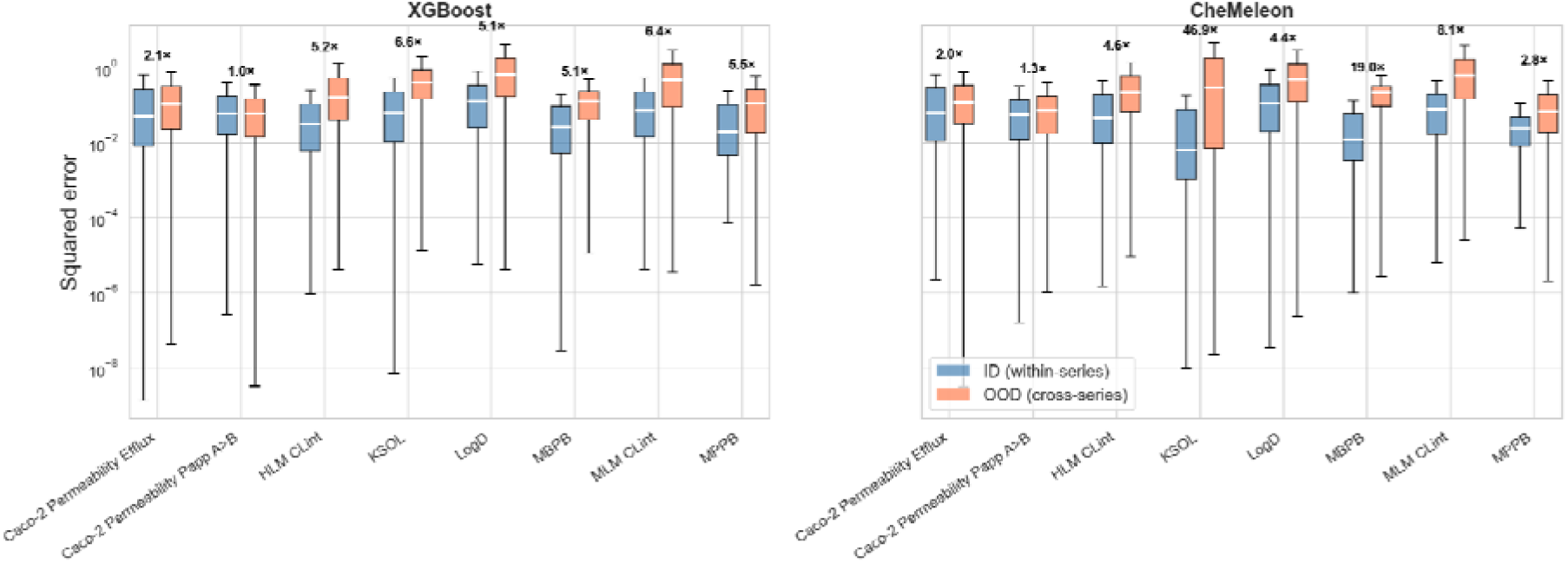
ID (within-series) versus OOD (cross-series) squared error distributions across 8 endpoints (MGMB excluded due to sparse coverage) for Optuna TPE-tuned XGBoost and CheMeleon foundation model (fine-tuned per endpoint). Box plots show median (white line), interquartile range, and whiskers (1.5 × IQR) on a log scale; outliers suppressed for clarity. Fold-change in median squared error annotated above each pair. R² goes negative OOD for 5/8 endpoints under XGBoost and 6/8 under CheMeleon.

**Table 4.**
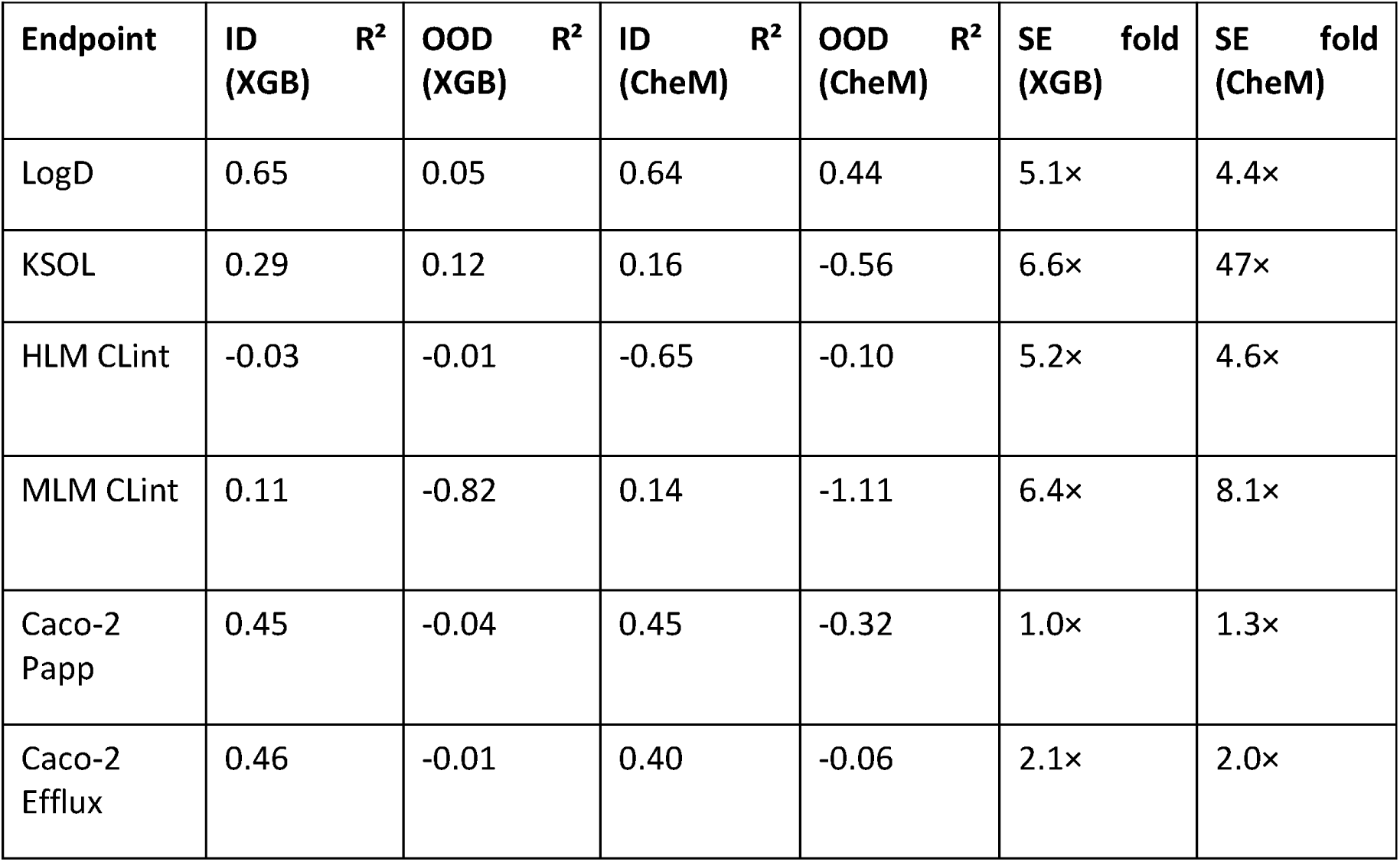
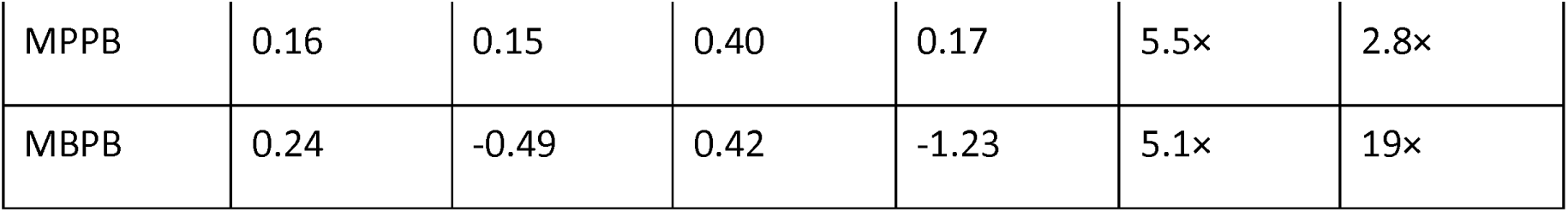
ID versus OOD performance across chemical series, XGBoost and CheMeleon results. ID denotes validation on 20% of the same largest Butina cluster used for training; OOD denotes testing on the second-largest Butina cluster, a structurally distinct chemical series absent from training. R² is the coefficient of determination on the corresponding held-out set (R² = 0 equals a constant mean-predictor baseline; R² < 0 is worse than that baseline). SE fold-change is the median squared error on OOD molecules divided by the median squared error on ID molecules, so values above 1 indicate larger typical per-molecule errors when crossing the chemical-series boundary.

The CheMeleon foundation model also does not close the ID–OOD performance gap. R² is negative on OOD for 6 of 8 endpoints (vs 5/8 under XGBoost): only LogD shows a meaningful OOD improvement (R² 0.44 vs 0.05 under XGBoost), while KSOL worsens (OOD R² −0.56), MBPB worsens (OOD R²-1.23), MLM CLint worsens (OOD R²-1.11), and Caco-2 Papp worsens (OOD R² −0.32). Here, foundation-model representation does not solve cross-series generalization: it improves in-distribution baselines while leaving (and in places amplifying) the structural cliff at series boundaries.

### Failure mode 2a: scaffold splits do not create meaningful distribution shift relative to random splits

Scaffold splits based on Bemis-Murcko decomposition are widely used as a proxy for evaluating model generalization on novel chemistry, under the assumption that holding out entire scaffolds creates a meaningful distribution shift.^31,32^ This assumption has been repeatedly challenged: distinct Bemis-Murcko scaffolds are often mutually similar, causing scaffold splits to retain high train-test similarity and overestimate performance relative to realistic settings.^33–35^ Consistent with this, we find that for this dataset, scaffold splitting confers no meaningful advantage over random splitting as an estimate of cross-series generalization. This result should not be read as a rejection of all series-aware splitting: alternative series definitions such as SCINS or medicinal-chemist-like automated series classification may be better aligned with some project questions.^36,37^ The point is that the split must be validated empirically. Distance-aware methods such as the cluster-based approach used here are necessary to ensure that test molecules are structurally distant from training molecules, for example measured with Morgan fingerprints, which are close proxy to the definition of molecular similarity in the context of medicinal chemistry, or by a separate design decision taken by the modelers.^11,38^

For both XGBoost and CheMeleon, we compared Bemis-Murcko scaffold splits to random and cluster-based splits and found that the common assumption, that scaffold-based holdout creates meaningful distribution shifts, does not hold for this dataset. Across 5-fold cross-validation on all 9 endpoints, Bemis-Murcko scaffold splits under XGBoost produced MA-RAE of 0.508, marginally worse than random splits (0.474) and more optimistic than cluster-based splits (0.674) (Figure S5, Table S2). Using CheMeleon, the corresponding values are 0.471 / 0.430 / 0.623, which are better than the XGBoost models but the relative ordering and gaps are preserved (scaffold–random ≈ 0.04, cluster–scaffold ≈ 0.15). The median test-to-train 1-NN distance for scaffold splits (0.246) is modestly larger than for random splits (0.203; two-sample KS D statistic = 0.20, p < 10⁻¹⁰), while cluster-based splits produce a substantially right-shifted distribution (median 0.424; KS D statistic = 0.66 vs random, p < 10⁻¹⁰). For this dataset, we find that scaffold splits produce nearly the same distances as random splits.

The failure of scaffold splits to create genuine distribution shifts has a structural explanation. Of 3,337 unique Bemis-Murcko scaffolds in this dataset, 2,252 (67.5%) contain only a single molecule, and the median scaffold group size is 1. We therefore argue that such greedy assignment of singleton scaffolds to folds is indistinguishable from random assignment. Furthermore, 56.4% of molecules (4,289 of 7,608) have their 1-NN in a different scaffold group, meaning that the scaffold boundary is structurally leaky, with cross-scaffold 1-NN distances (median 0.23) nearly overlapping overall 1-NN distances (median 0.19). Even among non-singleton molecules, 37.2% have a closer neighbor in a different scaffold group than within their own. We find that naive Bemis-Murcko scaffold splits do not necessarily provide a meaningful extrapolative test; in this dataset the large number of singleton scaffolds and the frequent crossing of nearest neighbors across scaffold boundaries make scaffold splits much closer to random splitting than is often assumed.

Importantly, Bemis-Murcko scaffolds carry biological signals beyond structural similarity alone. In a controlled comparison of molecule pairs at matched fingerprint distances, same-scaffold pairs exhibited significantly smaller activity differences than different-scaffold pairs (Mann-Whitney U test across 136 endpoint per distance-bin combinations; 91 significant after Benjamini-Hochberg correction at FDR < 0.05, of which 83 (91%) confirm that same-scaffold pairs have smaller |Δactivity|; Figure S6). This effect is strongest for HLM CLint (16/16 tested bins showing significantly smaller |Δactivity| for same-scaffold pairs), MLM CLint (14/17), and Caco-2 Efflux (12/15), consistent with endpoints where a protein must recognize the scaffold’s three-dimensional shape (CYP450 binding, P-glycoprotein substrate recognition). It is weakest for the protein binding endpoints MBPB (2/15) and MPPB (6/15). KSOL is the one endpoint where the direction is inconsistent: 10 of 17 bins reach significance but only 5 favour same-scaffold pairs, consistent with solubility being driven by substituent-level physicochemistry rather than scaffold identity.

However, the naive Bemis-Murcko scaffold split used here cannot leverage this biological signal. The split is fixed by the decomposition, leaving the user with no parameter to tune the degree of distribution shift to match a deployment scenario. The prevalence of singleton scaffolds (67.5% of scaffold groups) ensures most scaffolds are effectively randomly assigned, and the same rigid boundary is applied regardless of which endpoint is being predicted. Distance-based or explicitly series-aware methods give the user more direct control: clustering or series-definition parameters can be chosen explicitly, the grouping can be defined in a chosen structural feature space, there are no singleton artifacts by construction when groups are filtered or merged, and the degree of structural novelty can be explicitly matched to the deployment scenario.^11,39^

### Failure mode 2b: random cross-validation is precise at the wrong deployment level

Random folds are, by construction, unbiased draws from the same joint distribution over compounds and labels, so low between-repeat variance is expected. We analyzed whether the low variance of random cross-validation (CV) reflects genuine robustness or simply a stable artifact of information leakage. We compared the variance of 5 random 5-fold CV repeats to 5 cluster-based repeats. Across 5 independent random 5-fold CV repeats, RAE standard deviations were remarkably small (0.003–0.012), and ranges were tight (0.007–0.032) (Figure S7, Table S3). By contrast, 5 cluster-based repeats produced wider ranges on 8 of 9 endpoints (cluster/random std ratio: median 3.5×, spanning 0.6×–13.5×), with MGMB exhibiting the most extreme case (R² spanning 0.32–0.58 across repeats); KSOL was the only endpoint where the cluster spread was tighter than the random spread. A single cluster-split result could therefore report either poor or moderate model performance depending solely on the random seed.

Random 5-fold CV produces consistent performance but leak structurally similar molecules across fold boundaries (median test-to-train 1-NN of 0.20), creating an artificially easy evaluation that underestimates the challenges of predicting genuinely novel compounds, and is thus not relevant for structurally novel hit-identification or lead-identification scenarios. The gap between random (MA-RAE ∼0.47) and cluster-based (MA-RAE ∼0.67) evaluation, nearly 0.20 in MA-RAE, is roughly 7× the mean within-cluster per-endpoint split standard deviation and ∼33× the mean within-random per-endpoint split standard deviation, and the two distributions are non-overlapping (Mann-Whitney U = 25.0, p = 0.0079 for all 9 endpoints). The same Mann-Whitney result (U = 25, p = 0.0079, all 9 endpoints) holds under the CheMeleon foundation model. The choice of splitting strategy is therefore a far more consequential decision than the model architecture, number of repeats, or the hyperparameter search budget.

The practical recommendation is twofold: use distance-aware splits with multiple repeats, and always report confidence intervals. Five cluster repeats with wide CIs give a more honest picture than five random repeats with narrow CIs, because the former captures the genuine uncertainty about how a model will perform when deployed on novel chemistry within the diversity already represented in the dataset. This diagnostic is bounded by the training data itself: if the available dataset is narrow relative to the chemical space the model will ultimately be deployed against, splitting strategy alone cannot compensate for that gap, and cluster-based estimates may still be optimistic relative to true deployment performance in an unexplored region.

### Failure mode 3: activity cliffs expose interpolation failures

Activity or property cliffs carry high SAR informational content, as small structural changes associated with large potency shifts are often the most valuable cases for lead optimization^14^. Addressing this challenge likely requires either richer molecular representations capable of encoding the specific structural determinants driving activity discontinuities, or explicit modeling of structure-activity boundaries, directions reflected in activity-cliff benchmarking literature.^40^

To evaluate the impact of activity cliffs on prediction error, we first identified cliff pairs in advance based on structural similarity and activity difference criteria, then assessed model performance separately on cliff and non-cliff molecules. Using an absolute threshold of 1.0 log units (approximately 10-fold change in raw measurement), activity cliff molecules comprised 0–4.6% of compounds per endpoint, with LogD (4.6%), KSOL (3.5%), and MLM CLint (2.5%) showing the highest prevalence (Table S4). MBPB and MPPB had insufficient cliff molecules (0 and 4, respectively) for separate evaluation. Despite being a minority, these are typically of interest in practice since these molecules represent a qualitatively distinct failure mode from the extrapolation failures described earlier.

For both XGBoost and CheMeleon, we generated out-of-fold predictions under cluster-split cross-validation (5 folds) for all molecules and compared the error between cliff and non-cliff subsets (i.e. compounds that form or not form an activity cliff pair in the dataset). Cliff molecules showed systematically higher prediction error across all 7 endpoints with sufficient cliff populations in both models (Figure 7). Under XGBoost, the effects were most prominent for Caco-2 Efflux (cliff RAE 0.96 versus non-cliff 0.59), Caco-2 Papp A>B (cliff RAE 0.88 versus non-cliff 0.65), and MGMB (cliff RAE 0.78 versus non-cliff 0.55). Cliff R² is negative or near-zero for Caco-2 Efflux (−0.006) and MGMB (−0.23) under XGBoost; model performance on those cliff subsets is no better than predicting the cliff-set mean. These endpoint differences may be consistent with localized SAR effects, including transporter or metabolic-recognition motifs, but the result needed for the framework is empirical: molecules forming property cliffs are harder to predict under both architectures. CheMeleon also does not selectively help cliff molecules: cliff RAE still exceeds non-cliff RAE for all 7 endpoints, and although non-cliff RAE improves for 5 of 7 endpoints (largest drops: LogD 0.13, KSOL 0.08, MGMB 0.05), it is essentially unchanged for Caco-2 Efflux and worsens slightly for Caco-2 Papp A>B. The cliff/non-cliff RAE gap remains 0.06–0.44 (median ∼0.15). Foundation-model representation therefore can provide more accurate predictions for the non-cliff compounds but does not solve the relevant challenge of SAR discontinuities, and interpolation failures at activity cliffs occur for both models.

**Figure 7.**
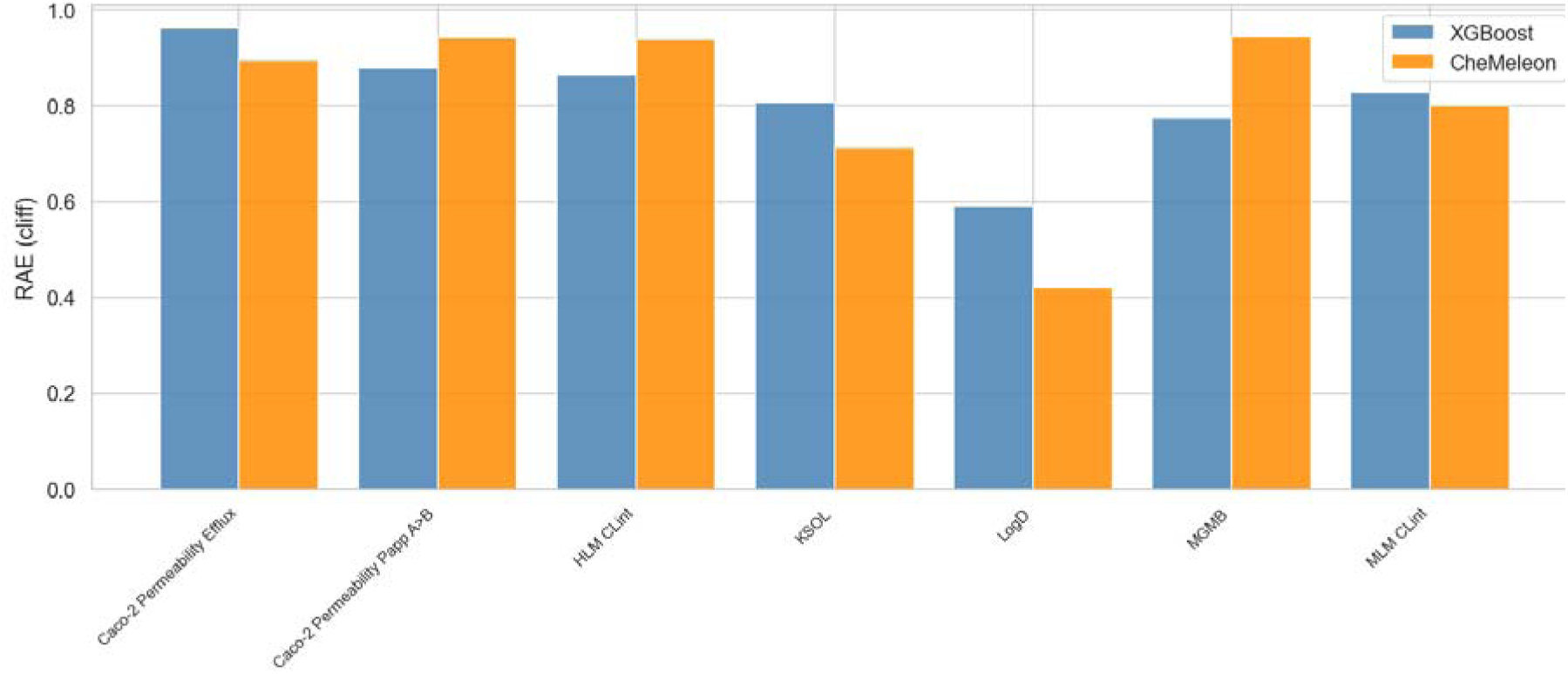
Prediction error (RAE) for cliff molecules only, by endpoint, comparing XGBoost and CheMeleon under cluster-split cross-validation. Cliff molecules participate in at least one pair with ECFP4 Tanimoto similarity ≥ 0.85 and absolute activity difference ≥ 1.0 log unit. Corresponding non-cliff RAE values are reported in Table S4.

The 0.85 Tanimoto threshold on ECFP4 is a common choice in ADME/QSAR practice but not a universal convention; the activity-cliff literature spans several operating points. Stumpfe & Bajorath (2014) identify cliffs as MMPs with ≥100× (≥2 log unit) potency difference, with their 0.85 threshold applying to MACCS keys (the ECFP4 value yielding the same similar-pair fraction is ∼0.56).^14^ MoleculeACE (van Tilborg et al. 2022) uses a soft consensus of ≥0.9 ECFP4, scaffold, or SMILES Levenshtein similarity together with ≥10× (≥1 log unit) potency.^15^ We therefore verified our 0.85 ECFP4 / ≥1 log unit definition against our data directly. Sweeping the similarity threshold across {0.70, 0.75, 0.80, 0.85, 0.90, 0.95} shows the RAE penalty emerges between 0.80 and 0.85. For similarity thresholds below 0.80 the cliff/non-cliff RAE gap is near-zero or negative for LogD, KSOL and MLM CLint, and at 0.90 three of nine endpoints have zero cliff pairs, increasing to four of nine at 0.95 (Figure S8).

### Failure mode 4a: molecular representations smooth stereoisomers and amplify scaffold decorations

Molecular property prediction models can only respond to differences that their input representation exposes. If a structural fingerprint assigns similar vectors to molecules with different biological properties, the model is blind to that difference, regardless of the training data or splitting strategy. This representation fidelity problem is particularly acute for stereoisomers, where enantiomers can differ in ADME properties due to stereoselective enzyme and transporter recognition, and for scaffold decorations where minor substituent changes can disproportionately alter the fingerprint vector. We therefore tested two controlled variant types: stereoisomers, where the 2D graph is identical but properties can differ, and scaffold decorations, where small substituent changes can create larger representation changes.

The extended Morgan fingerprints specification includes an option for encoding stereochemistry,^41^ but the extent of chirality sensitivity depends on the implementation. Enabling chirality-aware fingerprints (useChirality=True) in the RDKit allows the model to distinguish stereoisomers, but the signal remains weak (Figure S9, Table 5). Across 548 stereoisomer groups (1,152 molecules, 15.1% of the dataset, Table 5), chirality-aware Morgan fingerprints produce non-zero but small Tanimoto distances (median 0.07). The weakness has a concrete structural explanation: chirality-aware Morgan fingerprints encode stereochemistry by modifying the atom invariant at each stereocenter, but this modification only propagates through circular substructures that include that atom within the given radius. At radius 2 (ECFP4), only substructures within 2 bonds of the stereocenter are affected. Since 71% of stereoisomer molecules in this dataset have a single stereocenter, inverting the CW/CCW flag modifies a handful of substructure hashes, a median of 4 bits out of 2,048 (0.2%). 9% of stereoisomer pairs have zero bit flips and are completely invisible to the fingerprint (ECFP4 with radius 2), even with chirality enabled; a further 38% flip only 1–2 bits. Protonation preprocessing does not contribute to this blindness: dimorphite_dl preserves all stereocenters. Increasing the radius to 3 (ECFP6) doubles the median Tanimoto distance (0.14) and reduces the zero-flip fraction from 9% to 6%, confirming that radius is part of the bottleneck, though even at radius 3 the chirality signal remains small relative to the full fingerprint. Only 2 of 217 RDKit 2D descriptors differ between stereoisomers in this dataset (NumAtomStereoCenters and NumUnspecifiedAtomStereoCenters), contributing negligibly beyond the fingerprint signal. To quantify the downstream effect on predictions, we generated out-of-fold predictions for both XGBoost and CheMeleon under cluster-split cross-validation (5 folds) and computed the prediction coefficient of variation (standard deviation of predictions divided by the absolute mean) within each stereoisomer group. Under XGBoost, mean prediction coefficients of variation were 0.00–0.03 across all 9 endpoints, lower than scaffold decoration groups (mean CV 0.08–0.26) and random pairs (mean CV 0.13–0.38), indicating that the XGBoost model produces near-identical predictions for stereoisomers (Figure S10, Table S5, Figure S11).

**Table 5.**
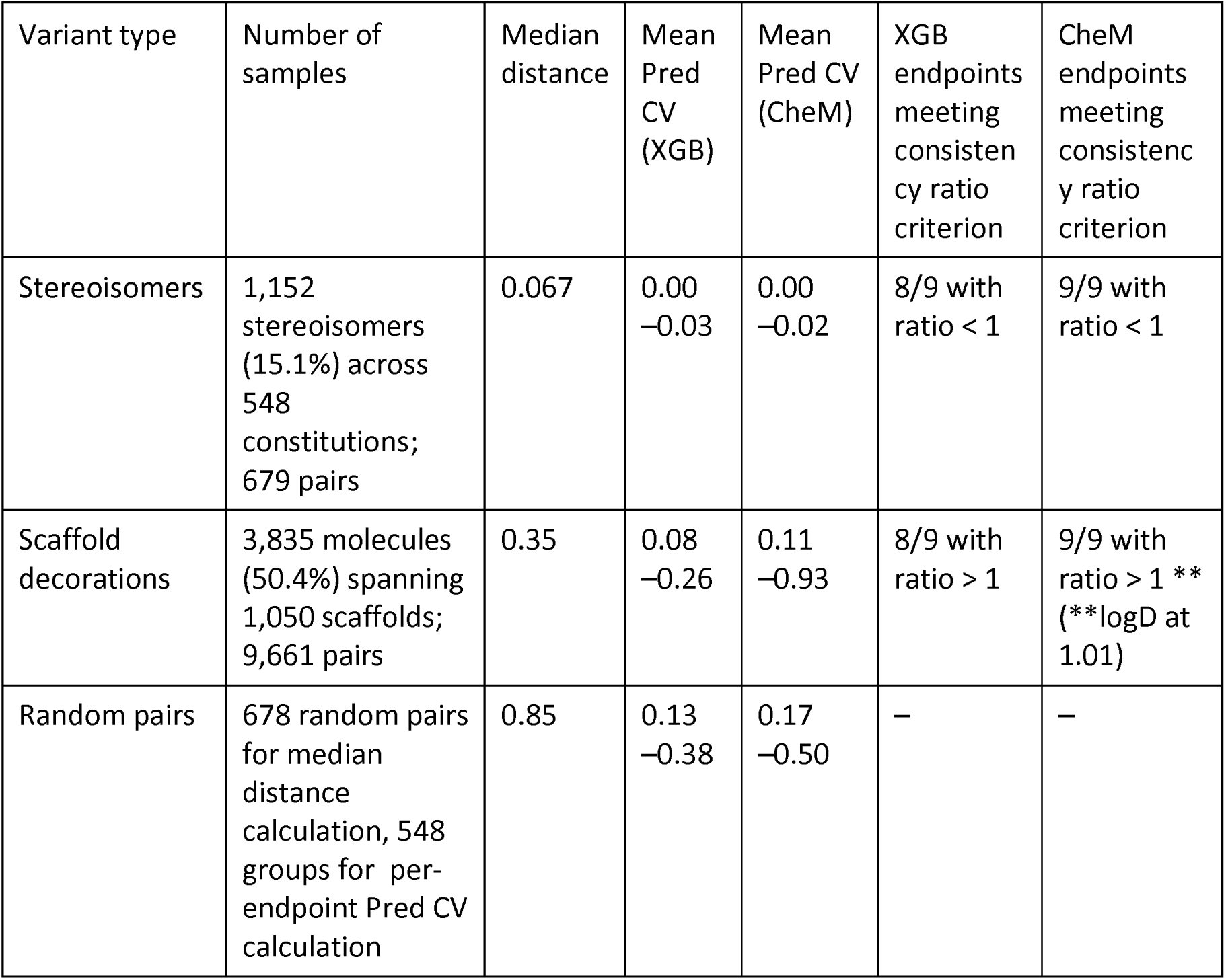
Prediction consistency by variant type, with side-by-side XGBoost and CheMeleon results. Prediction CV is the within-group standard deviation of predictions divided by the absolute group mean prediction. The consistency ratio is the within-group standard deviation of predictions divided by the within-group standard deviation of observed activities; ratios < 1 indicate under-prediction of observed variation, ratios > 1 indicate over-prediction. ‘Endpoints meeting criterion’ counts endpoints (of 9) with mean ratio on the stated side of 1.0.

It is known, however, that stereoisomers often do not show activities alike, including in ADME properties^42^. Enantiomers of the same compound may differ in metabolic clearance (stereoselective CYP450 oxidation, as with the warfarin enantiomers^43^), in plasma protein binding (chiral binding sites on serum albumin and alpha-1-acid glycoprotein^44^), and, for some substrates, in transporter-mediated efflux (stereoselective P-glycoprotein transport, which is substrate-dependent rather than universal^45^). In this dataset, stereoisomer pairs show non-trivial activity differences across endpoints (e.g., median |Δ| of 38 mL/min/kg for MLM CLint, 15 µM for KSOL, 0.10 units for LogD). However, an important caveat is that the dataset contains no repeat measurements of the same molecule, so the observed within-group variation reflects a mixture of true stereoselective effects and assay variability; these contributions cannot be fully separated without replicate data. Despite this ambiguity, mean consistency ratios (standard deviation of predictions divided by standard deviation of true values) remain below 1.0 for 8 of 9 stereoisomer endpoints (KSOL is the exception at 1.35), indicating the model under-predicts the total observed variation between stereoisomers for most endpoints, whether biological or experimental in origin. The chirality signal in the fingerprint is present but weak relative to the magnitude of within-group differences for 15% of the dataset. The CheMeleon model is somewhat more sensitive. Its architecture is based on Chemprop v2, whose featurization pipeline encodes an atom-level chiral tag at each stereocenter (derived from the RDKit’s GetChiralTag() function), giving the graph representation a signal for tetrahedral stereochemistry distinct from the substructure-hashing mechanism used by ECFP4. This raises the mean prediction CV to 0.00–0.02, but consistency ratios remain below 1.0 for all 9 endpoints under CheMeleon, including KSOL (0.72). This persistent under-prediction suggests that better methods for incorporating stereochemistry into 2D graph representations are needed.

Grouping molecules by their Bemis-Murcko scaffold and retaining groups of 2–20 members yielded 1,050 scaffold decoration groups containing 3,835 molecules (50.4% of the dataset, Table 5), each representing a set of analogs that share the same core but differ in their substituents. These groups revealed a complementary failure mode. They showed a median intra-group Tanimoto distance of 0.35 and prediction coefficients of variation lower than random pairs for 8 of 9 endpoints under both models (except LogD), confirming the models have learned meaningful scaffold-level trends. However, mean consistency ratios exceeded 1.0 for 8 of 9 endpoints under XGBoost (reaching 3.44 for KSOL, 2.70 for MGMB, and 2.04 for Caco-2 Papp A>B; LogD is the only endpoint with a ratio below 1) and for all 9 endpoints under CheMeleon (albeit logD with a ration at 1.01, see Table S5), indicating that both models may have over-predicted the substituent effects beyond true biological variation in a subset of groups; the ratio distribution is right-skewed, so the mean is driven by groups with small measured variation. Under XGBoost, this amplification is due to the discreteness of the Morgan representation: a single-atom decoration modifies the circular substructure containing that atom, flipping multiple bits. We find that the pretrained CheMeleon embedding is not measurably smoother than Morgan representation: on a scale-free measure (the percentile rank of a decoration partner among all 7,608 molecules, invariant to any monotone transform of the distance) the two representations are equivalent (median 0.145% vs 0.118%). The over-predicted substituent effects is therefore a shared consequence of predicting properties from local-substructure similarity.

These two failure modes, (1) under-prediction of stereochemical variation and (2) over-prediction of variation across scaffold decorations, stem from a disconnect between structure-representation distances and the biological distances that matter for ADME prediction. The representation imposes an implicit similarity function on molecules that may not align with the biological similarity function governing the endpoints of interest. While enabling chirality-aware fingerprints partially addresses the stereochemistry gap, 3D descriptors could further improve stereochemical sensitivity. This scaffold-decoration over-prediction appears in both circular fingerprints and graph inputs, indicating it is not specific to one representation. More broadly, the representation failures identified here highlight the complementary potential of bioactivity-based prediction approaches, such as transcriptomic signatures or Cell Painting morphological profiles, which encode biological response directly rather than relying on structural similarity, and may capture activity-relevant variation that fixed-radius fingerprints miss.

A practical caveat applies to chirality-aware modeling more broadly. Many public datasets (and likely, pharmaceutical datasets) contain unreliable stereochemistry annotations, arising from racemic synthesis, incomplete chiral separation, or inconsistent registration practices, which can make chirality-aware modeling counterproductive by fitting noise rather than signal. The decision to include or exclude chirality should therefore be informed by the reliability of the stereochemical annotations in the dataset at hand.

### Failure mode 4b: resonance form ambiguity causes asymmetric prediction risk

Molecules with delocalized electrons, such as those exhibiting multiple resonance forms, cannot be fully captured by a single Lewis structure, which forms the basis of the molecular representations used in property prediction models. Different but equally valid Lewis structures of the same molecule can therefore yield different molecular graphs, fingerprints, and descriptors, and consequently different model predictions. Following the evaluation framework of Zalte et al.(Zalte et al. 2025) who quantified this effect for molecular property prediction models, we assessed whether ADME predictions are stable under choice of resonance form using the pipeline described in the Methods section.

The fraction of molecules generating more than one distinct resonance form varies substantially by endpoint, from 19.6% (HLM CLint) to 53.4% (MGMB) at pH 7.4; at pH 6.5 (Caco-2 endpoints), only 1.8% do, as the lower pH protonates basic nitrogens and collapses most variants. Despite being chemically identical, resonance forms of the same molecule occupy distant regions of fingerprint space: median ECFP4 Tanimoto distance is 0.46, larger than scaffold decorations (0.35) and far larger than stereoisomers (0.067) (Figure 8A, Figure S12).

**Figure 8.**
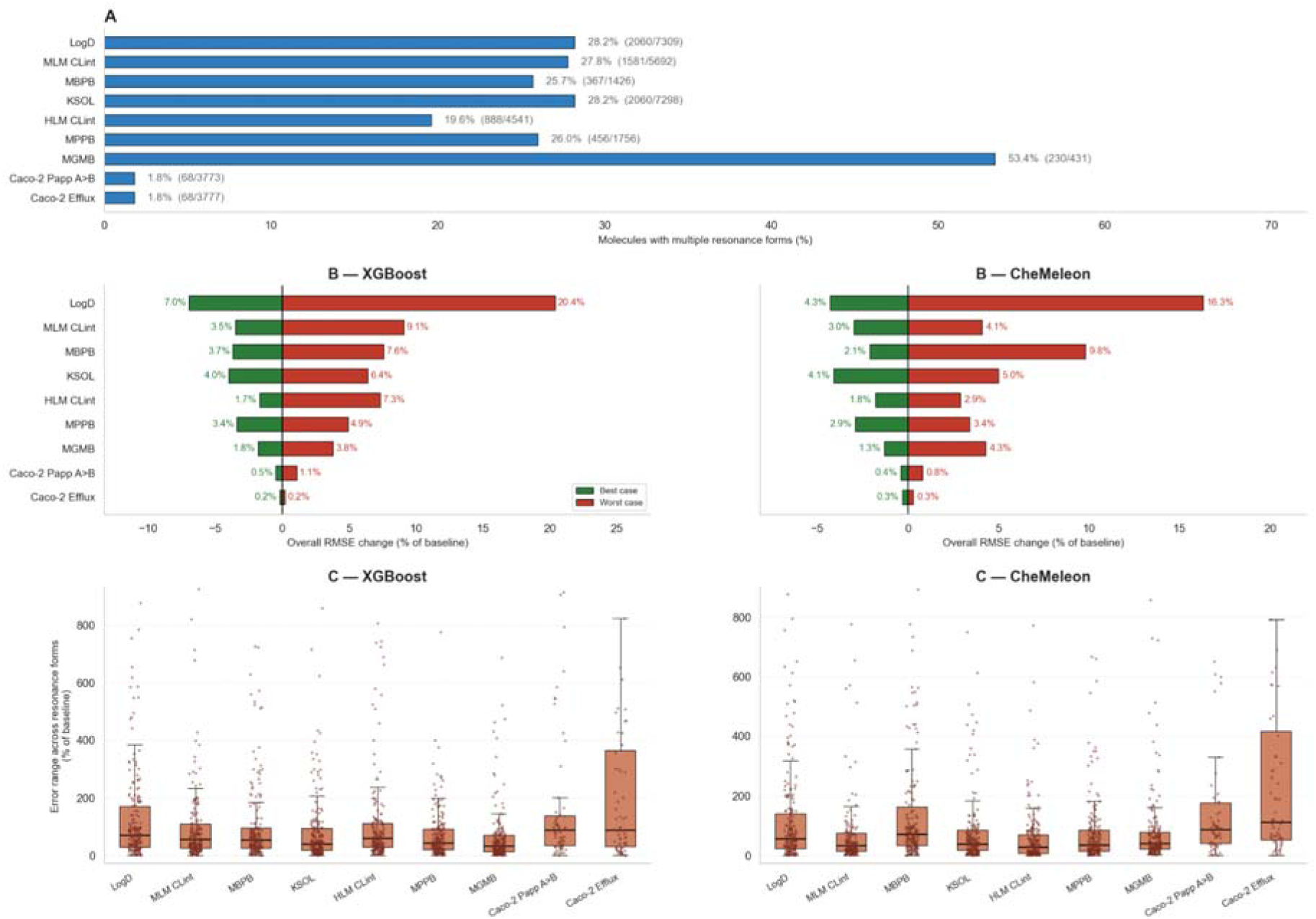
Sensitivity of ADME predictions to resonance form representation. (A) Fraction of molecules with more than one distinct resonance form per endpoint; pH 6.5 protonation (Caco-2) collapses most variants. (B) Change in RMSE, computed across all molecules for the endpoint, when the most favorable (green) or least favorable (red) resonance form is used as input, relative to baseline; worsening meets or exceeds improvement across all 9 endpoints in both models. Left: XGBoost. Right: CheMeleon. (C) Per-molecule prediction error range across resonance forms, normalized by baseline error, for molecules with >1 distinct form. Left: XGBoost. Right: CheMeleon.

For each molecule and endpoint, we compared predictions made on the input form (Baseline RMSE), the most favorable resonance form (RMS MinRD), and the least favorable (RMS MaxRD). Under XGBoost, total RMSE variation reaches 27.4% for LogD, with the worst-case form increasing RMSE by 20.4% and the best-case decreasing it by 7.0% (baseline RMSE 0.60). Material variation is observed across MLM CLint (12.6%), MBPB (11.3%), KSOL (10.3%), HLM CLint (9.0%), MPPB (8.3%), and MGMB (5.7%), while Caco-2 endpoints are minimally affected (≤1.5%) due to resonance collapse at pH 6.5 (Figure 8B, Table S6). Per-molecule prediction error ranges can exceed several hundred percent of baseline error (Figure 8C). The key concern is not the absolute magnitude but the invisibility of this sensitivity: standard evaluation pipelines commit to a single input form and report a single RMSE, with no indication of how much that number depends on an arbitrary representational choice.

CheMeleon is often, but not uniformly, more robust to resonance ambiguity than XGBoost, with lower total variation for LogD (20.6% vs 27.4%), MLM CLint (7.1% vs 12.6%), HLM CLint (4.7% vs 9.0%), and MPPB (6.3% vs 8.3%), comparable sensitivity for KSOL, MGMB, and Caco-2 Papp, and marginally higher for MBPB (11.9% vs 11.3%) and Caco-2 Efflux (0.5% vs 0.4%). That neither architecture fully escapes the effect is unsurprising: bond-order shifts and aromaticity changes propagate into whichever representation the model consumes, whether a molecular graph or a fingerprint vector. One proposed solution is to enforce invariance by construction by dropping the features that are sensitive to resonance form choice, as demonstrated for graph-based models by RIGR ^16^ Alternatively, augmenting training data with enumerated resonance forms can implicitly teach models to produce consistent predictions across representations. More broadly, resonance ambiguity is a data and representation issue that warrants explicit consideration when designing and evaluating models for chemical spaces with a high prevalence of delocalized systems.

## Discussion

We have presented a framework to evaluate generalization for molecular property prediction, and demonstrated its use on the Expansion Therapeutics ADMET dataset, a uniquely suited in-house dataset from real-world drug discovery programs. Each modeling case study is run under two architectures, an Optuna-tuned XGBoost with engineered features and a CheMeleon graph-foundation model fine-tuned per endpoint, so cross-model agreement isolates failure modes that originate in the data or representation. Through systematic case studies, we identified four complementary failure modes that standard aggregate metrics obscure. First, extrapolation failure: models degrade when crossing chemical space boundaries or target value ranges; R² goes negative on 5/8 endpoints under XGBoost (and 6/8 under CheMeleon) when training on one Butina cluster and testing on a different one. Second, interpolation failure: activity cliff molecules show systematically higher prediction error across all 7 endpoints with sufficient cliff populations, in both XGBoost and CheMeleon. Third, representation failure: chirality-aware Morgan fingerprints partially resolve stereoisomer blindness but still under-predict true biological variation (consistency ratio < 1 on 8/9 endpoints using XGBoost, and 9/9 using CheMeleon), while simultaneously over-predicting scaffold decoration effects beyond true biological variation; resonance form ambiguity causes RMSE to vary by 27.4% under XGBoost and 20.6% under CheMeleon, with worsening matching or exceeding improvement on 9/9 endpoints in both models. Fourth, evaluation failure: naive scaffold splits are indistinguishable from random splits (MA-RAE 0.51 vs 0.47), and random cross-validation produces precise metrics at an optimistically biased level (between-strategy gap ∼33× the mean within-random per-endpoint std). The foundation-model architecture lifts in-distribution baselines on 8/9 endpoints (from competition-split MA-RAE 0.78 to 0.68) but does not eliminate any of the failure modes; they are structural features of how chemical datasets are organized and how molecular representations encode them.

One route to improve performance is to augment chemical structure representation with a bioactivity-based one. For example, Cell Painting morphological profiles and L1000 transcriptomic signatures encode biological response directly, so structurally similar compounds with divergent activity are also separated in representation space.^46–48^ Empirically, Cell Painting–guided selection recovers activity-cliff pairs that structural similarity misses.^49^ This advantage is bounded by two practical constraints: profiles require the compound to be synthesized and assayed, placing them downstream of hit identification; and the reference panel (cell line, perturbation duration, feature set) defines a biological applicability domain that is typically narrower than the chemical space covered by ECFP4.^50,51^

Overall, findings in this work have direct practical implications. The choice of splitting strategy should be anchored to the deployment scenario (Table 1), and split choice matters far more than simply running more repeats: cluster-based splitting with confidence intervals provides more informative performance estimates for structural novelty, while random CV gives false confidence for that deployment scenario.^52,53^ Performance-over-distance curves should accompany single-number metrics, as they reveal where models fail and enable performance estimation for specific deployment sets. Molecular representations should be evaluated for blind spots (e.g., enabling chirality-aware fingerprints partially addresses stereoisomer blindness), but residual under-prediction of stereochemical variation, present in both ECFP4 and graph-foundation representations, suggests that 3D descriptors may be needed for full stereochemical sensitivity where reliable stereochemistry information is available.^54^ In the dataset, explored in this study, a graph foundation model improved baseline performance materially without changing any of the qualitative failure-mode conclusions.

While demonstrated on the Expansion Therapeutics dataset, these findings reflect principles that should apply broadly to molecular property prediction tasks with concentrated chemical series, sparse endpoint coverage, or deployment distributions that differ from training data. The four failure modes: extrapolation, interpolation, representation, and evaluation, are not artifacts of this particular dataset but arise from recognizable features of how molecular ML models learn and how real-world chemical datasets are organized. Datasets with medicinal-chemistry series are likely to exhibit extrapolation failure across chemical-space boundaries^55^; structurally local models can struggle at activity cliffs because cliffs create localized discontinuities in the structure-activity landscape;^56^ fixed structural representations have blind spots in their similarity functions; and evaluations that do not match the deployment distribution can produce misleadingly optimistic metrics. The recommendations (Box 1) are therefore meant as a reusable evaluation workflow, not as a claim that one data split or performance metric is universally sufficient.

## Methods

### Dataset

The Expansion Therapeutics ADMET dataset was obtained from the OpenADMET competition hosted on Hugging Face ^23,24^. The dataset comprises 7,608 ML-ready molecules from RNA-targeted small molecule drug discovery programs, with a predefined competition split (time split of 5,326 training and 2,282 test molecules with no overlap between the two sets) and nine ADME endpoints. The endpoints are: octanol-water distribution coefficient (LogD), kinetic solubility (KSOL, µM), human and mouse liver microsomal clearance (HLM CLint and MLM CLint, mL/min/kg), Caco-2 Apparent Permeability coefficient (Papp A>B, 10^-6^ cm/s), Caco-2 efflux ratio, and mouse plasma/microsomal protein binding (MPPB, MBPB, MGMB, expressed as percent unbound). Data were generated by four contract research organizations (Aragen, Chempartner, Pharmaron, WuXi) as well as through internal assays. Ordinal compound identifiers (formatted as E-XXXXXXX) preserve temporal ordering of synthesis, enabling time-based data splitting without requiring explicit timestamps.

### Molecular representations and feature computation

Both modeling approaches share the same preprocessing pipeline. Molecules were protonated at assay-relevant pH using dimorphite_dl:^57^ pH 7.4 for LogD, KSOL, HLM CLint, MLM CLint, MPPB, MBPB, and MGMB; pH 6.5 for Caco-2 Papp A>B and Caco-2 Efflux. The protonation window was set to pH ± 0.5, returning the single most probable protonation state. No salt or counterion stripping was required, as the dataset contains only single-fragment SMILES with no multi-component entries.

For the extreme gradient boosting (XGBoost) models, molecular fingerprints capturing atoms and local surroundings up to two bonds away were computed as Morgan fingerprints (radius 2, 2048 bits, and useChirality=True), functionally equivalent to the extended-connectivity fingerprint of diameter 4 (ECFP4)^41^, using the RDKit ^58^. Physicochemical descriptors were computed using the full RDKit 2D descriptor suite (approximately 200 descriptors). Zero-variance RDKit 2D descriptors were removed, while fingerprint bits were retained without variance filtering, because XGBoost handles sparse binary features natively^59^., Descriptors were normalized using StandardScaler (scikit-learn ^60^), fit on training data and applied to test data. The final feature vector for each molecule was the concatenation of the 2048-bit Morgan fingerprint and the scaled RDKit 2D descriptors, yielding approximately 2,200 features for the XGBoost model. Target values for all endpoints except LogD were logarithmically transformed using the formula log10(clip(x, 10^-10) + 1), matching the competition evaluation protocol.

### Structural distance computation

Pairwise structural distances were computed as Jaccard distances (1 minus Tanimoto similarity) between 2048-bit Morgan fingerprints. For each test molecule, the distance to the nearest training neighbor (1-NN) was used as the primary measure of structural novelty. Chemical diversity was assessed by computing all-pairwise, 1-nearest-neighbor (1-NN), and 5-nearest-neighbor Jaccard distance distributions. Chemical series were identified using Butina clustering ^61^ with a Jaccard distance cutoff of 0.7.

The Jaccard distance of Morgan fingerprints was chosen as the primary distance metric for various reasons. First, similarity thresholds for this metric have been empirically characterized across hundreds of activity classes, making distance thresholds more interpretable and comparable across studies.^39,62^ Second, ECFP4 Tanimoto coefficients (equivalent to Morgan fingerprints and Jaccard index) are among the most widely used similarity metrics in cheminformatics, providing a shared reference frame for the community.^63,64^ Third, the distance metric operates in the same feature space as the XGBoost fingerprint input and provides a fixed structural-novelty axis for comparing both model classes. Fourth, a permissive Tanimoto similarity > 0.3 for Morgan fingerprints captures most co-active compounds compared to similarity between random compounds, providing empirically grounded thresholds for defining structural novelty.^39^ Usually compounds with Tanimoto similarity > 0.85 have been considered structurally similar compounds ^34,65^, but we acknowledge that Morgan fingerprint Tanimoto similarity has known limitations: it is insensitive to three-dimensional molecular shape, biased toward scaffold identity over substituent effects, and cannot capture pharmacophoric similarity between topologically distinct molecules. These limitations motivate the complementary analyses (molecular variants, activity cliffs) presented in this work.

### Cross-validation splitting strategies

Three splitting strategies were implemented, each designed to mimic a different deployment scenario: cluster-based splitting (structural generalization), time-based splitting (temporal generalization), and target property-value splitting (value extrapolation).

For cluster-based splitting, molecules were embedded into a Euclidean space via an Empirical Kernel Map: kernel principal component analysis (PCA) was applied to the precomputed Tanimoto similarity matrix and the leading 50 components retained.^60^ Mini-batch k-means was then applied in this projected space to produce 20 clusters using three random initializations and a batch size capped at 1,024 molecules which were greedily assigned to five folds by repeatedly placing the largest remaining cluster into the fold holding the fewest molecules.^60^ This over-clustering strategy ensures balanced fold sizes (max/min ratio < 1.15 across all endpoints). Fold assignments were computed per endpoint, using only molecules with non-null values. Five independent repeats were generated using different random seeds (0–4), yielding 25 train/test splits per endpoint. For time-based splitting (temporal generalization), molecules were sorted by their ordinal index and divided into five equal groups. We employed an expanding-window cross-validation scheme across four folds to evaluate model performance on successive ‘future’ data. Specifically, for each fold k the model was trained on the union of all preceding cohorts and evaluated on the subsequent unseen cohort. For target-value splitting (value extrapolation), molecules were sorted by their endpoint value and divided into five equal groups, using the same expanding-window scheme. In both time-based and target-value strategies, the first group serves exclusively as the initial training set and is never used as a test fold, yielding 4 test folds from 5 groups.

Splitting strategies were validated using a set of quantitative diagnostics to ensure robustness and consistency across folds. Fold size balance was assessed by verifying that the ratio between the largest and smallest test folds remained below 1.15 across all endpoints and strategies. Target distributions were evaluated by comparing per-fold endpoint value distributions using Kolmogorov Smirnov (KS) tests, enabling quantification of train–test distributional shifts within each fold. Structural separation was examined through test-to-train NN distance analysis, where the median Jaccard distance from each test molecule to the training set was computed per fold, confirming that each strategy achieved its intended level of structural dissimilarity. Structural overlap was further quantified as the proportion of test molecules having a training neighbor within a Jaccard distance threshold of 0.1, with cluster-based splitting approaches exhibiting minimal to negligible overlap. Finally, Uniform Manifold Approximation and Projection (UMAP) embeddings derived from a precomputed Jaccard distance matrix were used to visualize the data, demonstrating that fold assignments preserved cluster integrity as well as temporal and value-based ordering.

Adversarial validation, where a classifier is trained to distinguish training from test molecules, can provide a complementary diagnostic of split severity^66^; To quantify covariate shift with a model-based criterion, we applied adversarial validation to each splitting strategy and was run on LogD (7,309 molecules) as a representative endpoint. For every fold, test molecules were labelled 1 and their training molecules 0, and a gradient-boosted classifier (HistGradientBoostingClassifier) was trained on 2048-bit ECFP4 fingerprints to distinguish them, scored by 3-fold cross-validated AUROC and averaged across folds.

### Model training and evaluation

Every modeling case study was run under two architectures so that failures attributable to data and evaluation design can be distinguished from representation-specific effects. The first architecture, XGBoost^59^ used Optuna TPE (Tree-structured Parzen Estimator) hyperparameter optimization, ensuring that all analyses use per-endpoint tuned models rather than arbitrary default configurations. For each (endpoint, split strategy, fold) combination, nine hyperparameters were tuned using 30 Optuna trials with 3-fold inner cross-validation (TPE sampler, seed = 42): n_estimators (100–1000), max_depth (3–12), learning_rate (0.01–0.3, log-uniform), subsample (0.5–1.0), colsample_bytree (0.3–1.0), min_child_weight (1–10), gamma (0.0–5.0), reg_alpha (0.0–1.0), reg_lambda (0.5–3.0). Best parameters were cached per (endpoint, split strategy, fold) and reused across analyses sharing identical training sets, ensuring reproducibility and eliminating tuning as a source of variation between analyses. The second architecture, CheMeleon^67,68^, is a graph-based molecular foundation model fine-tuned per endpoint. Pre-trained weights for the GNN backbone were loaded and fine-tuned with a two-layer feed-forward head for each endpoint. Training used a 90/10 train/validation split with early stopping based on validation loss, with protonated SMILES as the sole input. Together, the two architectures bracket the modeling space, a tuned tree ensemble on engineered features versus a fine-tuned graph foundation model, and are evaluated on identical splits, endpoints, and protonated molecular forms with architecture-specific inputs, so cross-model agreement can isolate failure modes that originate in the data or evaluation rather than in the learning algorithm.

Model performance was assessed using mean absolute error (MAE; sklearn.metrics.mean_absolute_error), coefficient of determination (R^2; sklearn.metrics.r2_score), Spearman rank correlation (scipy.stats.spearmanr), Kendall rank correlation (scipy.stats.kendalltau), and relative absolute error (RAE = MAE / mean absolute deviation). We report R^2 rather than R^2_0 (the coefficient of determination for regression through the origin, a stricter external validation metric proposed by Golbraikh and Tropsha^69^) because the focus of this work is on comparing splitting strategies and failure modes rather than on external predictivity validation; all metrics are computed on held-out test folds under each splitting strategy. Because R^2 depends on the variance of the held-out target distribution, the ID-vs-OOD analysis also reports median squared-error fold-change, which captures the change in typical per-molecule error when crossing a chemical-series boundary. MA-RAE (macro-averaged RAE across 9 endpoints) served as the primary aggregate metric, as it normalizes for endpoint-specific variance and is interpretable as the fraction of baseline (mean-predictor) error retained by the model. Because MA-RAE weights all endpoints equally regardless of sample size, estimates for sparse endpoints carry greater fold-to-fold variance. For cluster-and time-based splits, coverage-weighted averaging changes MA-RAE by about 2% or less (cluster −0.8%, time −2.2%), so equal weighting does not materially change the aggregate metric; for target-value splits, coverage weighting increases MA-RAE materially because KSOL, the endpoint with the worst performing target-split RAE, also has the highest coverage, so up-weighting it dominates the aggregate. Thus, per-endpoint metrics are reported throughout to sidestep this aggregation sensitivity.

### Performance-over-distance curves

Both architectures were evaluated under all three splitting strategies (cluster-based with 5 folds, time-based with 4 folds, and target-value with 4 folds). Distances were binned using a sliding window: range defined as Q1 - 1.5 × IQR (interquartile range) to Q3 + 1.5 × IQR, bin width = (range)/5, step = (bin width)/20, minimum 25 molecules per bin. RMSE was computed per bin with median and 10th–90th percentile confidence bands across folds.

### In-distribution (ID) versus out-of-distribution (OOD) evaluation

For both XGBoost and CheMeleon, the largest Butina cluster (n = 2,572) was split temporally 80/20 for ID training (n = 2,057) and validation (n = 515). The second-largest cluster (n = 1,301) served as the OOD test set, representing a structurally distinct chemical series not seen during training.

To compare scaffold splits to random and cluster splits, both XGBoost and CheMeleon were evaluated against naive Bemis-Murcko^70^ scaffold splits generated by greedy assignment of scaffold groups to 5 folds, against random splits, and against cluster-based splits to test whether scaffold grouping creates meaningful distribution shift. To investigate the relationship between scaffold membership and structural separation, we computed scaffold boundary violation rates (fraction of molecules whose overall 1-NN has a different Bemis-Murcko scaffold) and cross-scaffold vs. within-scaffold proximity distributions. To assess whether scaffold membership carries biological information beyond fingerprint distance, we tested activity concordance conditioned on scaffold membership (Mann-Whitney U tests comparing |Δ_activity_| for same-scaffold vs. different-scaffold pairs within Tanimoto distance bins of width 0.05, across all 9 endpoints).

### Split variance analysis

For both architectures, we evaluated 5 random and 5 cluster-based 5-fold cross-validation repeats (seeds 0–4) across all 9 endpoints (∼450 fitted models per architecture, that is 10 repeats × 5 folds × 9 endpoints) to quantify within-strategy variance (RAE/R² spread across repeats) and between-strategy bias (random vs cluster). Per-endpoint cluster-repeat versus random-repeat RAE was compared with a two-sided Mann-Whitney U test.

### Activity cliff evaluation

Cliff pairs were defined as molecule pairs with Tanimoto similarity > 0.85 (Morgan fingerprints) and an absolute activity difference ≥ 1.0 log unit (for log-transformed endpoints, computed in log₁₀(x + 1) space, corresponding to approximately 10-fold change in raw measurement; for LogD, computed in raw units). Any molecule participating in at least one cliff pair was labeled a cliff molecule. After cliffs in the dataset were identified, their impact on performance was evaluated by comparing cliff versus non-cliff prediction error for each model (XGBoost and CheMeleon) under cluster-split CV

### Sensitivity to Stereochemical and scaffold decorations

Both XGBoost and CheMeleon were evaluated under cluster-split cross-validation (5 folds) for whether the models produce consistent predictions for molecules that differ in controlled ways. Two types of molecular variant groups were defined. Stereoisomer groups were constructed by stripping stereochemistry from each SMILES and grouping molecules that share the same achiral canonical SMILES. This yields pairs or small sets of enantiomers and diastereomers whose biological activity may differ substantially despite identical 2D structure. Scaffold decoration groups were constructed by computing Bemis-Murcko scaffolds and grouping molecules that share the same scaffold, retaining groups of size 2–20 to focus on series with meaningful but bounded structural variation.

For each group, we computed two metrics across each model’s predictions for group members: (1) prediction coefficient of variation (standard deviation of predictions divided by the absolute mean), which measures how much predictions vary within a group relative to their magnitude; and (2) consistency ratio (standard deviation of predictions divided by standard deviation of true values), which measures whether the model amplifies (ratio > 1) or smooths (ratio < 1) the true endpoint variation within the group. As a baseline, we constructed random groups by sampling molecules without replacement at sizes matched to the observed stereoisomer group sizes, and computed the same metrics.

### Resonance form sensitivity evaluation

Zalte et al.^16^ highlighted that different resonance forms of the same molecule can yield inconsistent predictions in graph-based models. To quantify the impact of this ambiguity on ADME predictions, resonance structures were enumerated for each molecule from the protonated SMILES using the RDKit’s native resonance enumeration function, capped at 50 structures per molecule.

Both XGBoost and CheMeleon were evaluated under cluster-split cross-validation to quantify the effect of resonance form ambiguity on model performance. For each test molecule, all distinct resonance forms were enumerated and predicted independently. Zalte et al.^16^ introduced RMS MaxRD as a worst-case measure of resonance-induced prediction variance, selecting the resonance form that maximizes deviation from the ground truth. Building on this, we additionally report the best-case analogue (RMS MinRD) and the total variation (RMS MaxRD − RMS MinRD). Together, these three metrics characterize the full range of ambiguity introduced by resonance form choice. To further characterize the structural diversity among resonance forms, Jaccard distances were computed between all pairwise resonance forms within each molecular group at pH 7.4, benchmarked against a size-matched sample of random molecule pairs.

## Conclusion

Adopting this framework will require a cultural shift beyond individual researchers. As a scientific community, we are interested in developing and improving methods in general as well as in making a contribution to drug discovery and a difference for patients. The latter requires a mindset that is different from textbook model building and validation. Journal and conference reviewers can create strong incentives by requiring evaluation against distribution-relevant splits, not just random splits, and by asking authors to characterize the type of distribution shift their model will face in deployment. In our companion work on method comparison best practices^7^, we argued for structured reporting and visual summaries; here, we extend that argument to model evaluation itself. The community has the tools; what is needed now is the expectation that rigorous generalization evaluation is the standard, not the exception. We hope this framework, together with the openly available Expansion Therapeutics dataset, encourages the community to move toward evaluation practices that better reflect the challenges of deploying molecular property predictions in real drug discovery.

## Code and Data Availability

All analysis code, notebooks, datasets and configuration files are freely available at https://github.com/srijitseal/polaris. The Expansion Therapeutics ADMET dataset is available at https://huggingface.co/datasets/openadmet/openadmet-expansionrx-challenge-data.

## Acknowledgements

S. Seal acknowledges support from awards from the NIH Complement Animal Research in Experimentation Promoting non-animal methods Challenge and the NIH Complement Animal Research in Experimentation NAMs Reduction to Practice Challenge. The authors would like to thank Pat Walters for his comments on an earlier version of the manuscript.

