## Supplementary Information for "Model Validation Protocols for Machine Learning in Small Molecule Drug Discovery"

1. Broad Institute of MIT and Harvard, Cambridge, MA, US.
2. Human Chemical Company, CA, US.
3. Uppsala University, Uppsala, Sweden.
4. Centre for Molecular Informatics, Yusuf Hamied Department of Chemistry, University of Cambridge, Lensfield Road, Cambridge CB2 1EW, United Kingdom
5. Department of Chemical Engineering, MIT, Cambridge, MA, US
6. Groningen Research Institute of Pharmacy, Groningen, NL
7. Department of Epidemiology, School of Public Health, University of São Paulo, Av. Dr. Arnaldo, 715, São Paulo, SP, 01246-904, Brazil
8. Novo Nordisk, Molecular AI, Lexington, MA, US.
9. Genentech, South San Francisco, CA, United States of America
10. UCB, Slough SL1 3WE, United Kingdom
11. College of Health and Wellbeing, University of Glasgow, Glasgow, United Kingdom
12. Lilly Research Laboratories, Eli Lilly and Company, Indianapolis, Indiana 46285, United States
13. Merck Healthcare KGaA, Darmstadt, Germany
14. Boehringer Ingelheim Pharma GmbH & Co. KG, Biberach, Germany.
15. Independent
16. Merck & Co., Inc., South San Francisco, California 94080, United States
17. Merck & Co., Inc., Cambridge, Massachusetts 02141, United States
18. Nimbus Therapeutics, Boston, MA, US.
19. Bayer Research and Innovation Center, Boston, MA, US.
20. Novartis Biomedical Research, Novartis Campus, 4002 Basel, Switzerland
21. Pfizer, Machine Learning Research, Berlin, Germany.
22. AstraZeneca R&D, Molecular AI, Gothenburg, Sweden.
23. Department of Computer Science and Engineering, Chalmers University of Technology and University of Gothenburg, Gothenburg, Sweden
24. Toxicology Data Science, Bayer SAS Crop Science Division, Sophia-Antipolis, Valbonne, France
25. F. Hoffmann-La Roche Ltd., Roche Innovation Center Basel, Switzerland.
26. Boehringer Ingelheim Pharma, London, UK
27. Expansion Therapeutics, Boston, MA, USA
28. Department of Medicine and Center for Biotechnology, College of Medicine and Health Sciences, Khalifa University of Science and Technology, Abu Dhabi, United Arab Emirates
29. STAR-UBB Institute, Babeş-Bolyai University, Cluj-Napoca, Romania
30. Research Center for Functional Genomics, Biomedicine and Translational Medicine, Iuliu Hatieganu University of Medicine and Pharmacy, Cluj-Napoca, Romania"
31. Valence Labs, Montreal, Canada
32. Recursion Pharmaceuticals, Montreal, Canada

† These authors contributed equally.

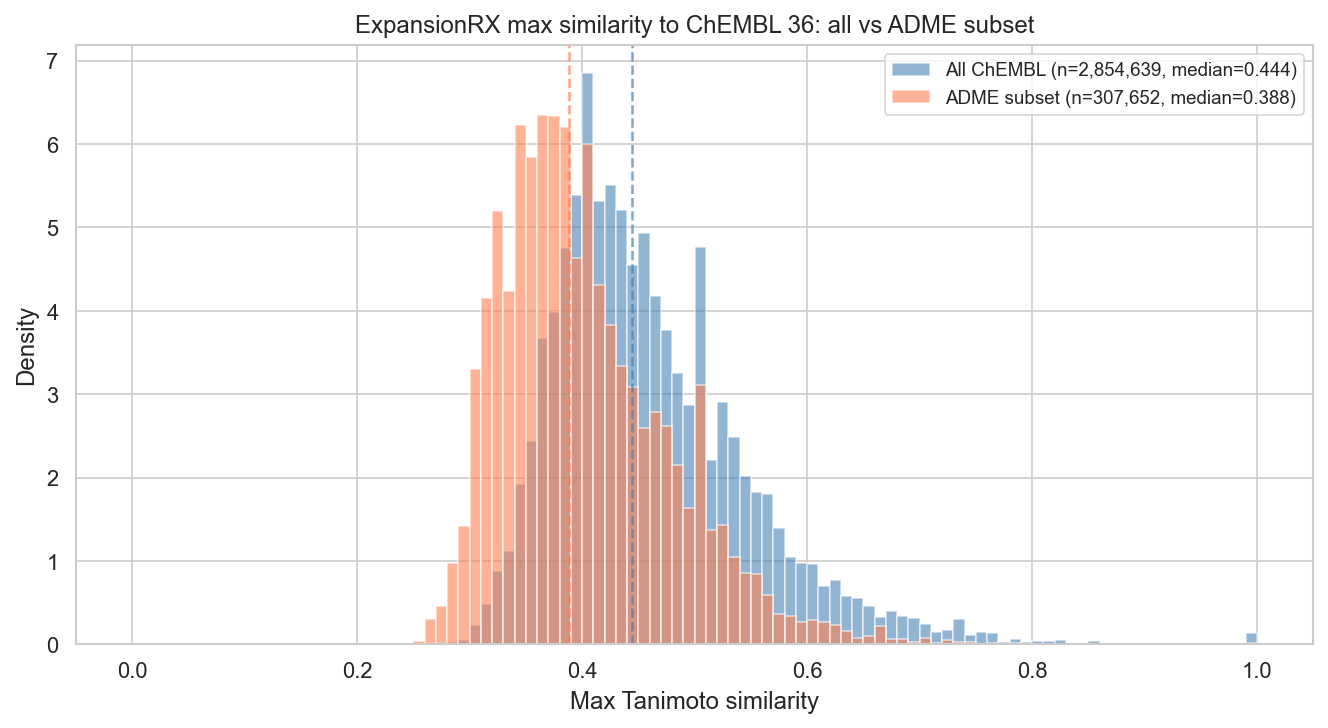

**Figure S1.** Maximum Tanimoto similarity of Expansion Therapeutics compounds to ChEMBL 36. Distribution of the highest Morgan fingerprint (radius 2, 2048-bit) Tanimoto similarity between each dataset molecule (n = 7,608) and (a) all ChEMBL 36 compounds (2.85M molecules, blue) or (b) the ADME-annotated subset (308K molecules, orange). Vertical dashed lines indicate medians (0.44 and 0.39, respectively). The 0.7 activity-relevant similarity threshold (dotted line) encompasses 98% of dataset compounds against all ChEMBL, indicating that most molecules in the Expansion Therapeutics dataset lack close structural analogues in the public domain.

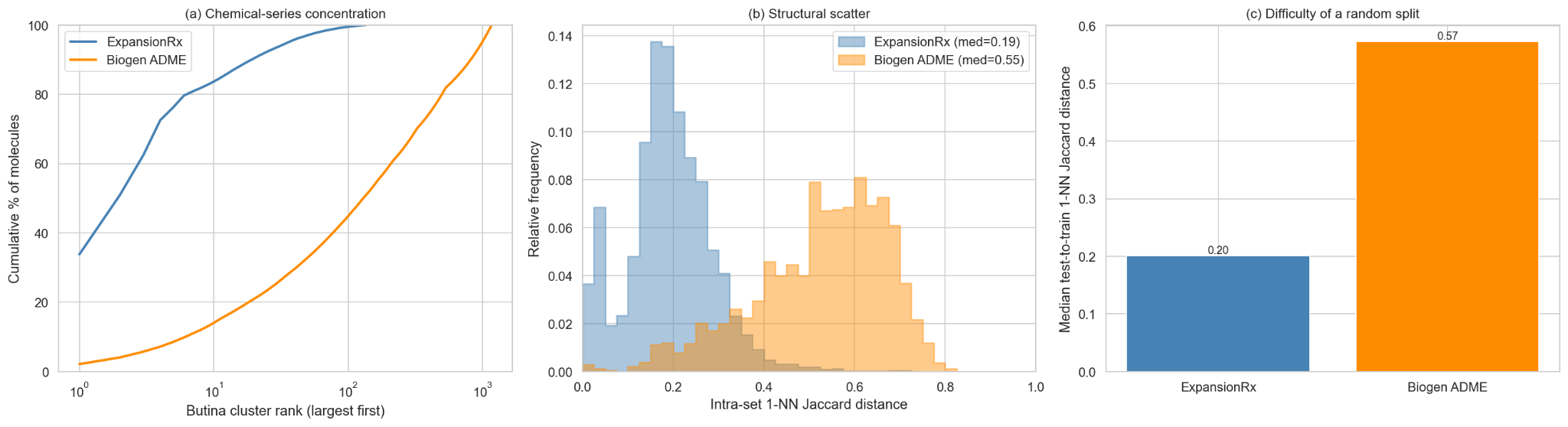

**Figure S2.** Dataset characterization of the ExpansionRx dataset (which is more concentrated in chemical series) and the public Biogen ADME dataset (which is more structurally global). All molecules in both datasets are represented by 2048-bit ECFP4 fingerprints (radius 2, chirality-aware) and compared by Tanimoto (1 − similarity = Jaccard) distance; clusters are defined by Butina clustering at a 0.7 distance cutoff. (a) Cumulative fraction of molecules covered as a function of Butina cluster rank (largest first, log scale). ExpansionRx concentrates into a few large series: its largest cluster holds 33.8% of compounds and the top ten cover 83.7% (135 clusters total), while Biogen has hundreds of clusters (1,169 total; largest 2.1%, top ten 14.0%, 54% singletons). (b) Distribution of each molecule's nearest-neighbour (1-NN) Jaccard distance to any other molecule in the same dataset (relative frequency); Biogen is shifted higher (median 0.55 vs 0.19), confirming its compounds are structurally dispersed rather than grouped into series. (c) Median test-to-train 1-NN Jaccard distance under a random 80/20 split, averaged over five random seeds. In Biogen a random split already places test molecules far from training (0.57), essentially equal to the dataset's intrinsic scatter, whereas in ExpansionRx, random test molecules remain close to training (0.20). Together these show that a series-based ID-vs-OOD split is not as such constructible in Biogen and that no "easy" random split exists there, illustrating that the appropriate splitting strategy is a property of the dataset, established by characterization, rather than a protocol applied uniformly.

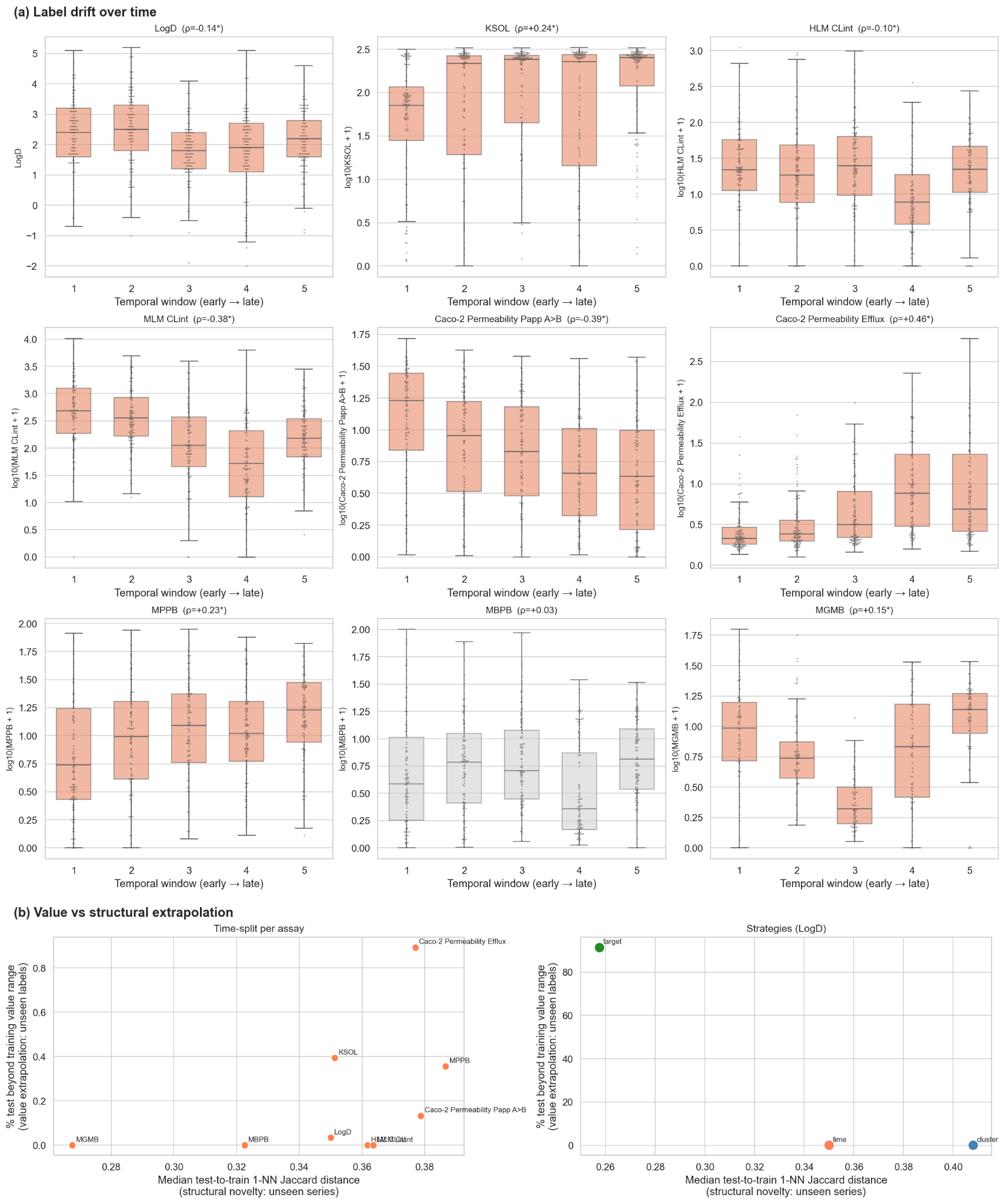

**Figure S3.** Temporal label drift across ADME assays; time-split tests structural novelty rather than value extrapolation. (a) Label drift over time. For each of the nine ADME assays, measured values are shown across the five temporal windows of the time-based split (window 1 = earliest compounds, window 5 = latest), where windows are the expanding-window folds defined by the ordinal compound index (a registration-order proxy for time). Each panel shows a box summarising every molecule in the window (median, interquartile range, 1.5 × interquartile range (IQR) whiskers; outliers hidden) with a swarm of a random subsample (≤120 points per window) overlaid to convey the underlying distribution: the swarm is illustrative and does not show every molecule. Values are log-transformed [log₁₀(value + 1)] except LogD, which is already on a log scale. Panel titles give the Spearman rank correlation (ρ) between compound index and value across the full assay; assays with appreciable drift (p < 0.05 and |ρ| ≥ 0.10) are coloured coral, the one assay without a reliable trend (mouse blood protein binding, MBPB) is grey. Eight of nine assays drift, and is strongest for the Caco-2 permeability assays, kinetic solubility (KSOL), and mouse liver microsomal clearance (MLM CLint). Rather, the direction is assay-specific: LogD and clearance trend down, solubility and efflux trend up. (b) Value versus structural extrapolation. Each point is placed by structural novelty (x-axis: median distance from each test molecule to its nearest training molecule, 1 − Tanimoto on ECFP4 fingerprints; higher = test chemistry less like training) and value extrapolation (y-axis: percentage of test molecules whose measured value falls beyond the training value range; higher = labels never seen in training). Left, the time-split for each assay; right, the three splitting strategies for LogD. The two axes are distinct: cluster-split maximises structural novelty while keeping all test values inside the seen range, the target-value split structurally closest to training yet 91.5% of test values beyond range, and is the only split that tests value extrapolation. The time-split lies between them on structure while inducing almost no value extrapolation (≤0.9% across all assays).

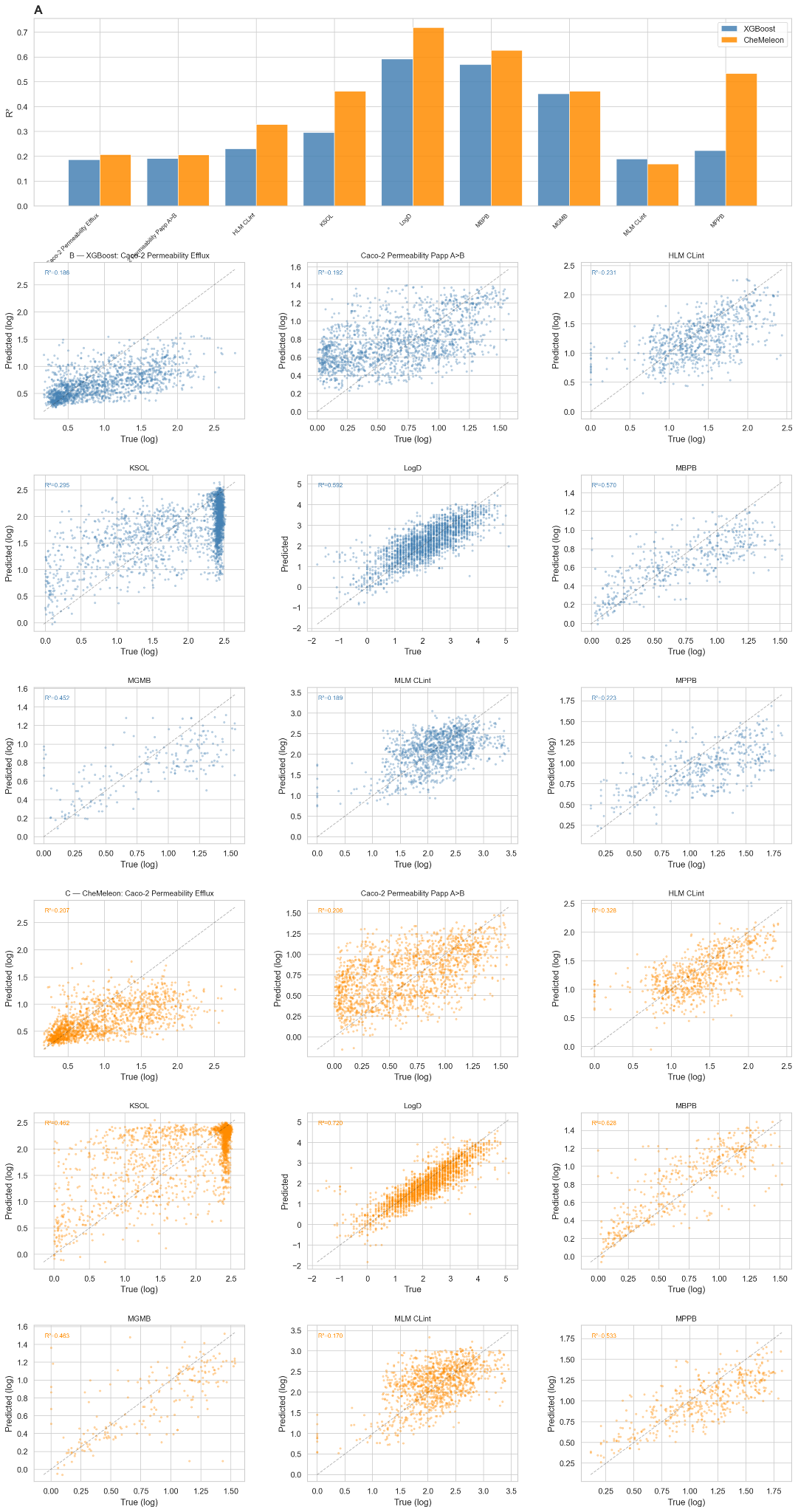

**Figure S4.** Baseline performance on the competition-split test set. (A) Per-endpoint R² comparing Optuna TPE-tuned XGBoost (Morgan fingerprints + RDKit 2D) with the CheMeleon graph-foundation model. CheMeleon improves R² on 8 of 9 endpoints; only MLM CLint regresses slightly (-0.02). Largest gains: MPPB (+0.31), KSOL (+0.17), and LogD (+0.13). (B) Predicted versus true values for XGBoost across all 9 endpoints; diagonal line indicates perfect prediction. (C) Predicted versus true values for CheMeleon.

**Table S1.** Baseline performance on the competition-split test set with side-by-side XGBoost and CheMeleon results. R² is the coefficient of determination on the held-out test set; Spearman is rank correlation between predicted and measured endpoint values; RAE is MAE divided by endpoint-specific mean absolute deviation; MA-RAE is the unweighted mean RAE across endpoints.

| **Endpoint** | **R² XGB** | **Spearman XGB** | **RAE XGB** | **R² CheM** | **Spearman CheM** | **RAE CheM** | **n_train** | **n_test** |
| --- | --- | --- | --- | --- | --- | --- | --- | --- |
| LogD | 0.59 | 0.76 | 0.63 | 0.72 | 0.87 | 0.49 | 5,039 | 2,270 |
| KSOL | 0.30 | 0.50 | 0.83 | 0.46 | 0.54 | 0.59 | 5,128 | 2,170 |
| HLM CLint | 0.23 | 0.56 | 0.85 | 0.33 | 0.63 | 0.80 | 3,759 | 782 |
| MLM CLint | 0.19 | 0.46 | 0.92 | 0.17 | 0.47 | 0.94 | 4,522 | 1,170 |
| Caco-2 Papp A>B | 0.19 | 0.51 | 0.86 | 0.21 | 0.54 | 0.83 | 2,157 | 1,616 |
| Caco-2 Efflux | 0.19 | 0.65 | 0.78 | 0.21 | 0.68 | 0.75 | 2,161 | 1,616 |
| MPPB | 0.22 | 0.62 | 0.88 | 0.53 | 0.75 | 0.67 | 1,302 | 454 |
| MBPB | 0.57 | 0.78 | 0.58 | 0.63 | 0.83 | 0.50 | 975 | 451 |
| MGMB | 0.45 | 0.69 | 0.66 | 0.46 | 0.72 | 0.58 | 222 | 209 |
| **MA-RAE** |  |  | **0.78** |  |  | **0.68** |  |  |

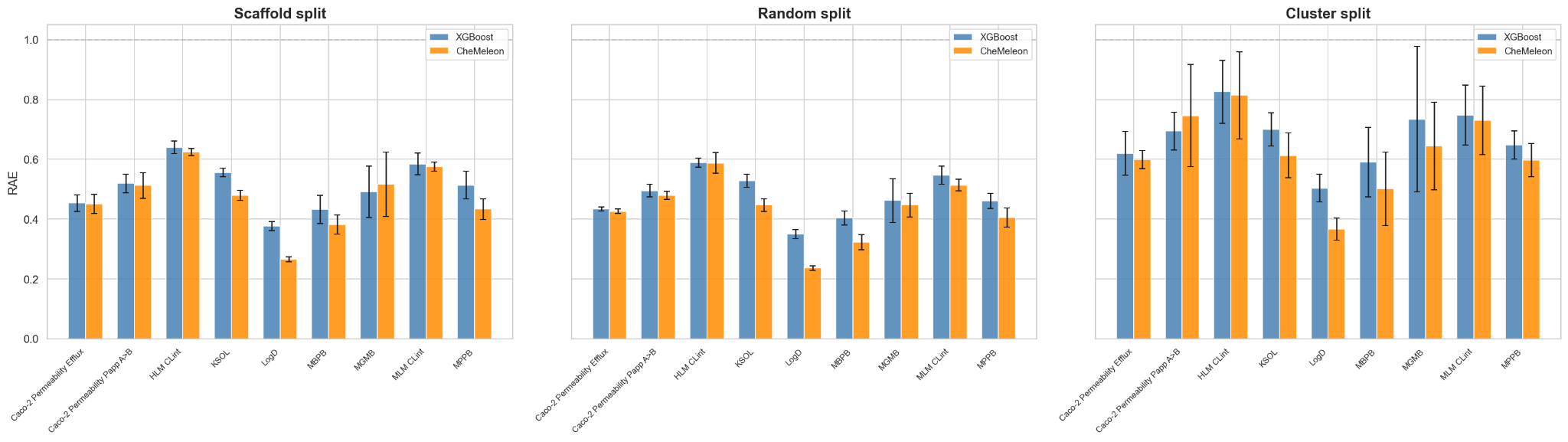

**Figure S5.** RAE by endpoint under three splitting strategies, comparing XGBoost and CheMeleon. Each panel shows one splitting strategy: scaffold (left), random (center), cluster (right), with XGBoost and CheMeleon bars side-by-side per endpoint. Error bars show standard deviation across 5 folds. RAE is relative absolute error normalized by the endpoint-specific mean absolute deviation; lower values are better, and RAE = 1 (dashed line) corresponds to the constant mean-predictor baseline. Across both model classes, scaffold splits remain close to random splits and substantially more optimistic than cluster splits.

**Table S2.** Per-endpoint RAE by splitting strategy with side-by-side XGBoost and CheMeleon results. RAE is MAE divided by endpoint-specific mean absolute deviation; values below 1 improve over the constant mean-predictor baseline. The scaffold-≈-random ≪ cluster ordering is preserved in both models.

| **Endpoint** | **Random XGB** | **Random CheM** | **Scaffold XGB** | **Scaffold CheM** | **Cluster XGB** | **Cluster CheM** |
| --- | --- | --- | --- | --- | --- | --- |
| LogD | 0.35 | 0.24 | 0.38 | 0.27 | 0.50 | 0.37 |
| KSOL | 0.53 | 0.45 | 0.56 | 0.48 | 0.70 | 0.61 |
| HLM CLint | 0.59 | 0.59 | 0.64 | 0.62 | 0.83 | 0.81 |
| MLM CLint | 0.55 | 0.51 | 0.58 | 0.57 | 0.75 | 0.73 |
| Caco-2 Papp A>B | 0.50 | 0.48 | 0.52 | 0.51 | 0.69 | 0.75 |
| Caco-2 Efflux | 0.43 | 0.43 | 0.45 | 0.45 | 0.62 | 0.60 |
| MPPB | 0.46 | 0.41 | 0.51 | 0.43 | 0.65 | 0.60 |
| MBPB | 0.40 | 0.32 | 0.43 | 0.38 | 0.59 | 0.50 |
| MGMB | 0.46 | 0.45 | 0.49 | 0.52 | 0.73 | 0.64 |
| **MA-RAE** | **0.47** | **0.43** | **0.51** | **0.47** | **0.67** | **0.62** |

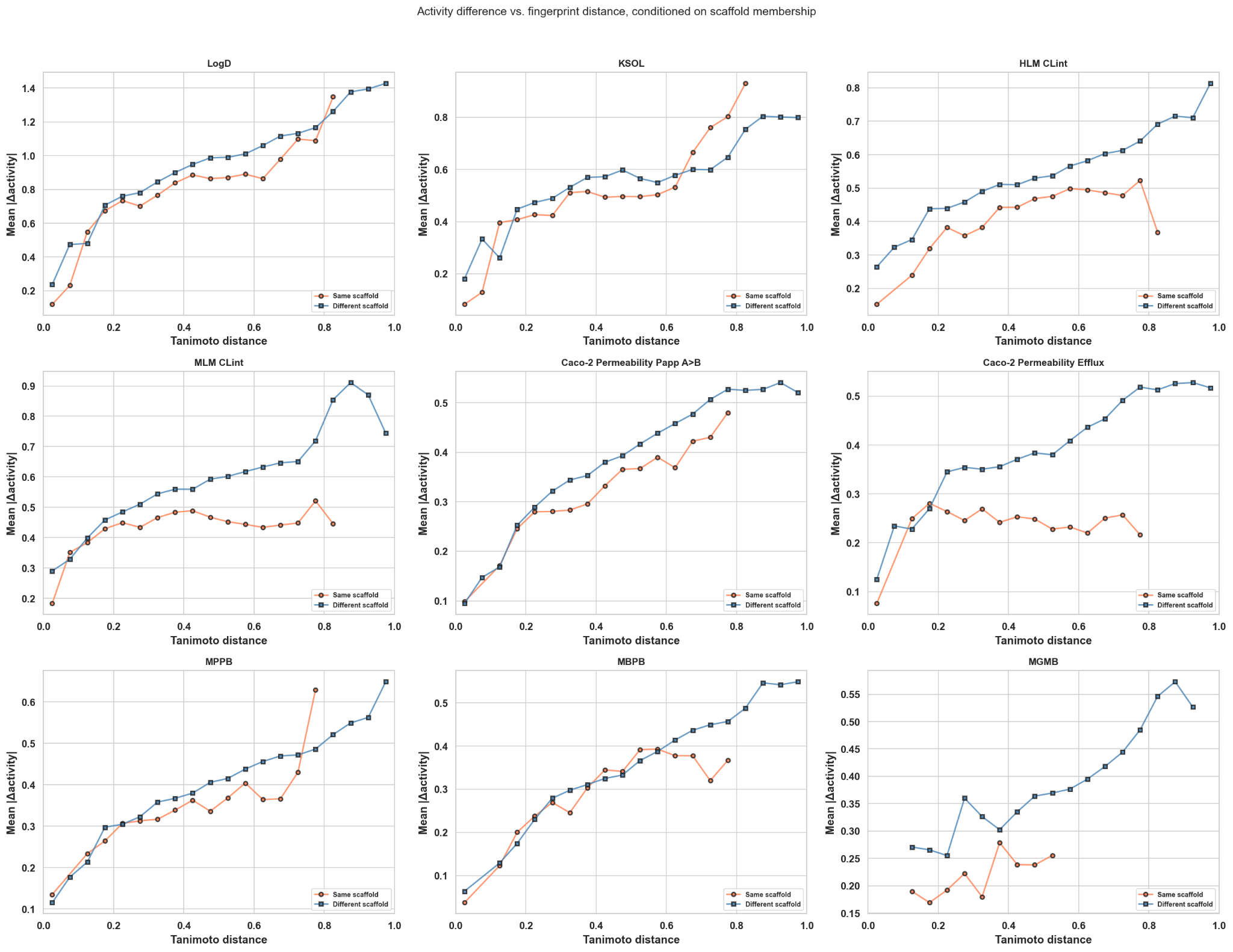

**Figure S6.** Mean pairwise activity difference (|y_i_ - y_j_|, where *y* is the activity value and *i* and *j* are two compounds) as a function of Jaccard distance (Morgan fingerprints, useChirality=True), stratified by scaffold membership (same vs. different Murcko framework), for all 9 endpoints. At a given fingerprint distance, same-scaffold pairs tend to have smaller activity differences, but the magnitude and consistency of this effect is highly endpoint-specific. The effect is strongest for Caco-2 Efflux and microsomal clearance, and weakest for protein binding endpoints.

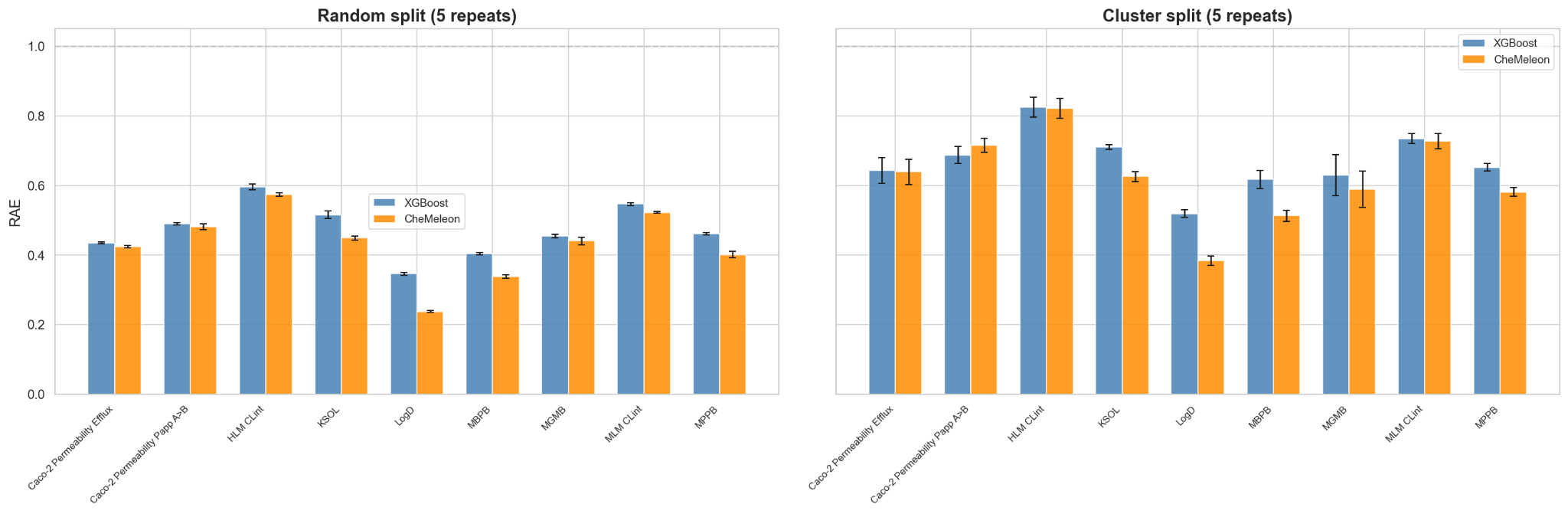

**Figure S7.** Random- and cluster-split RAE across 5 repeats, comparing XGBoost and CheMeleon. Each panel shows one splitting strategy: random (left) and cluster (right), with XGBoost and CheMeleon bars side-by-side per endpoint. Error bars show standard deviation across 5 independent repeats. Random splits produce tight variance at an optimistic level in both models; cluster splits reveal larger uncertainty at a harder level. The Mann-Whitney U test comparing cluster-repeat RAE to random-repeat RAE returns U = 25, p = 0.0079, the maximum possible separation for n₁ = n₂ = 5 — for all 9 endpoints in both XGBoost and CheMeleon.

**Table S3.** Performance variance across repeated splits, with side-by-side XGBoost and CheMeleon results. Entries are mean R² ± standard deviation across 5 independent repeats, with the minimum and maximum repeat values in parentheses. Random repeats and cluster repeats each use 5-fold cross-validation.

| **Endpoint** | **Random R² (XGB)** | **Random R² (CheM)** | **Cluster R² (XGB)** | **Cluster R² (CheM)** |
| --- | --- | --- | --- | --- |
| LogD | 0.86 ± 0.00 (0.86–0.87) | 0.92 ± 0.00 (0.92–0.93) | 0.70 ± 0.01 (0.69–0.72) | 0.82 ± 0.01 (0.81–0.83) |
| KSOL | 0.65 ± 0.01 (0.64–0.66) | 0.67 ± 0.00 (0.67–0.68) | 0.42 ± 0.01 (0.42–0.43) | 0.44 ± 0.02 (0.42–0.46) |
| HLM CLint | 0.60 ± 0.01 (0.58–0.62) | 0.62 ± 0.00 (0.62–0.63) | 0.27 ± 0.04 (0.22–0.34) | 0.26 ± 0.04 (0.21–0.33) |
| MLM CLint | 0.67 ± 0.00 (0.67–0.68) | 0.69 ± 0.00 (0.68–0.69) | 0.42 ± 0.03 (0.39–0.47) | 0.43 ± 0.04 (0.39–0.51) |
| Caco-2 Papp A>B | 0.70 ± 0.01 (0.69–0.70) | 0.68 ± 0.01 (0.67–0.70) | 0.45 ± 0.03 (0.42–0.49) | 0.37 ± 0.02 (0.33–0.39) |
| Caco-2 Efflux | 0.75 ± 0.00 (0.74–0.75) | 0.73 ± 0.01 (0.72–0.74) | 0.51 ± 0.04 (0.46–0.58) | 0.47 ± 0.06 (0.39–0.54) |
| MPPB | 0.74 ± 0.00 (0.74–0.74) | 0.79 ± 0.01 (0.78–0.80) | 0.53 ± 0.02 (0.50–0.54) | 0.59 ± 0.01 (0.58–0.61) |
| MBPB | 0.78 ± 0.00 (0.77–0.78) | 0.82 ± 0.00 (0.82–0.83) | 0.55 ± 0.03 (0.50–0.60) | 0.66 ± 0.02 (0.62–0.68) |
| MGMB | 0.69 ± 0.00 (0.69–0.70) | 0.68 ± 0.01 (0.65–0.69) | 0.48 ± 0.09 (0.32–0.58) | 0.47 ± 0.06 (0.39–0.57) |

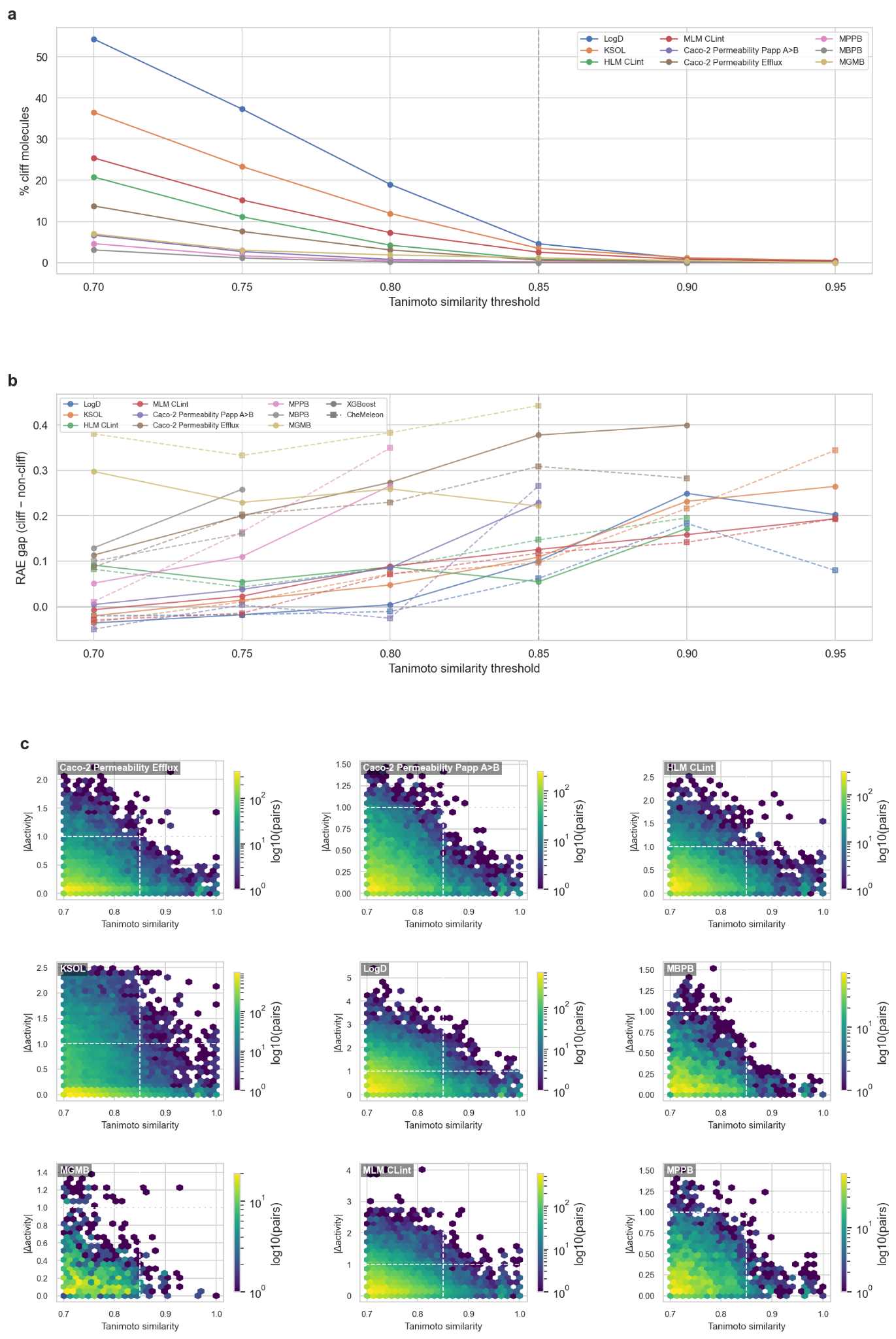

**Figure S8.** Sensitivity of the activity-cliff definition to the Tanimoto similarity threshold. The similarity cutoff is swept across {0.70, 0.75, 0.80, 0.85, 0.90, 0.95} on ECFP4 Tanimoto while the activity-difference criterion is held fixed at |Δactivity| ≥ 1.0 log unit. (a) Percentage of test molecules involved in at least one cliff pair, per endpoint. Mean prevalence falls ~12.7× between 0.70 and 0.85 and ~10.2× between 0.85 and 0.95; 3 of 9 endpoints reach zero cliff pairs at 0.90, rising to 4 of 9 at 0.95. Prevalence depends only on structural similarity and activity differences and is therefore model-independent. (b) RAE gap (cliff RAE - non-cliff RAE) per endpoint for XGBoost (solid circles) and CheMeleon (dashed squares); positive values indicate that cliff molecules are harder than non-cliff molecules. Below 0.80 the gap is near-zero or negative for LogD, KSOL, and MLM CLint in both models (the cliff label is not selective in this regime); above 0.90 the statistic becomes unstable as cliff populations collapse. The gap pattern is preserved between XGBoost and CheMeleon, interpolation failure at activity cliffs is not eliminated by the foundation-model representation. (c) Joint density (hexbin, log-scaled counts) of ECFP4 Tanimoto similarity versus |Δactivity| over all similar pairs (sim ≥ 0.70), one panel per endpoint. White dashed lines mark the adopted cliff definition at similarity = 0.85 and |Δactivity| = 1.0. Gray dashed verticals in (a) and (b) mark the same reference threshold.

**Table S4.** Activity cliff prevalence and performance impact under cluster-split cross-validation, with side-by-side XGBoost and CheMeleon RAE. Cliff molecules participate in at least one pair with ECFP4 Tanimoto similarity ≥ 0.85 and absolute activity difference ≥ 1.0 log unit. Percent cliff is the fraction of measured molecules for that endpoint labeled as cliff molecules. Dashes indicate too few cliff molecules for a stable cliff-specific RAE estimate; MPPB had only 4 cliff molecules and MBPB had 0 cliff pairs.

| **Endpoint** | **% cliff** | **Non-cliff RAE XGB** | **Cliff RAE XGB** | **Non-cliff RAE CheM** | **Cliff RAE CheM** |
| --- | --- | --- | --- | --- | --- |
| Caco-2 Efflux | 0.58% | 0.58 | 0.96 | 0.59 | 0.89 |
| Caco-2 Papp A>B | 0.16% | 0.65 | 0.88 | 0.68 | 0.94 |
| HLM CLint | 0.81% | 0.81 | 0.86 | 0.79 | 0.94 |
| MLM CLint | 2.5% | 0.70 | 0.83 | 0.68 | 0.80 |
| KSOL | 3.5% | 0.70 | 0.81 | 0.62 | 0.71 |
| LogD | 4.6% | 0.49 | 0.59 | 0.36 | 0.42 |
| MGMB | 1.2% | 0.55 | 0.78 | 0.50 | 0.94 |
| MPPB | 0.2% | 0.62 | – | 0.56 | – |
| MBPB | 0.0% | 0.56 | – | 0.48 | – |

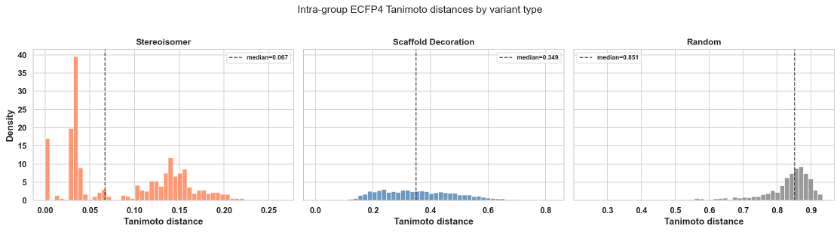

**Figure S9.** Intra-group Morgan fingerprints Jaccard distance distributions for stereoisomers (median 0.07), scaffold decorations (median 0.35), and random pairs (median 0.85). Chirality-aware fingerprints partially resolve stereoisomer blindness, but distances remain an order of magnitude smaller than those of scaffold decorations.

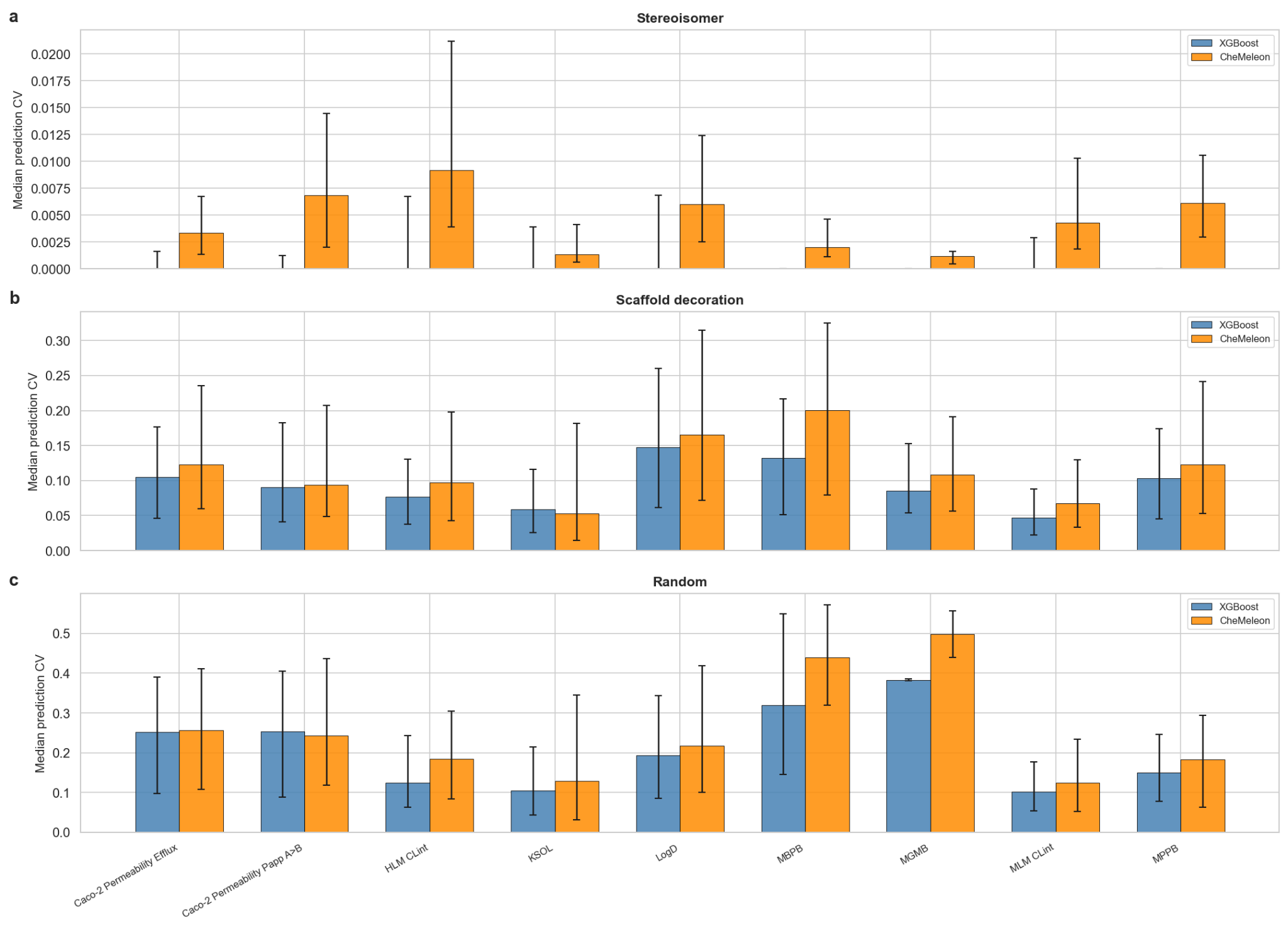

**Figure S10.** Per-endpoint prediction coefficient of variation for three variant group types, comparing XGBoost (blue) and CheMeleon (orange). Bars show the median prediction CV across groups; whiskers span the interquartile range (Q25–Q75). (a) Stereoisomer groups: median CV ≈ 0.00–0.01, confirming near-identical predictions for enantiomers under XGBoost (chirality-aware ECFP4) and only slightly higher under CheMeleon (graph representation registers stereo bond direction directly). (b) Scaffold decoration groups: intermediate CV (0.05–0.20), indicating models have learned scaffold-level trends but under-differentiate substituent effects. (c) Random pairs: upper bound CV (0.10–0.50). Note the y-axis scale differs across rows to preserve visibility of the small stereoisomer signal.

**Table S5.** Prediction coefficient of variation and consistency ratio by variant type and endpoint, with side-by-side XGBoost and CheMeleon results. Prediction CV measures how much predictions vary within a group relative to their magnitude. Consistency ratio measures whether the model amplifies (ratio > 1) or smooths (ratio < 1) the true within-group variation.

| **Variant type** | **Endpoint** | **Groups** | **Pred CV median (XGB)** | **Pred CV median (CheM)** | **Consistency ratio mean (XGB)** | **Consistency ratio mean (CheM)** |
| --- | --- | --- | --- | --- | --- | --- |
| Stereoisomer | LogD | 536 | 0.000 | 0.006 | 0.16 | 0.23 |
| Stereoisomer | KSOL | 536 | 0.000 | 0.001 | 1.35 | 0.72 |
| Stereoisomer | HLM CLint | 319 | 0.000 | 0.009 | 0.53 | 0.86 |
| Stereoisomer | MLM CLint | 394 | 0.000 | 0.004 | 0.37 | 0.56 |
| Stereoisomer | Caco-2 Papp A>B | 263 | 0.000 | 0.007 | 0.36 | 0.97 |
| Stereoisomer | Caco-2 Efflux | 263 | 0.000 | 0.003 | 0.19 | 0.29 |
| Stereoisomer | MPPB | 76 | 0.000 | 0.006 | 0.01 | 0.47 |
| Stereoisomer | MBPB | 67 | 0.000 | 0.002 | 0.00 | 0.52 |
| Stereoisomer | MGMB | 12 | 0.000 | 0.001 | 0.00 | 0.22 |
| Scaffold decoration | LogD | 1,007 | 0.148 | 0.165 | 0.92 | 1.01 |
| Scaffold decoration | KSOL | 999 | 0.059 | 0.053 | 3.44 | 2.92 |
| Scaffold decoration | HLM CLint | 635 | 0.077 | 0.097 | 1.23 | 1.65 |
| Scaffold decoration | MLM CLint | 762 | 0.047 | 0.068 | 1.21 | 1.66 |
| Scaffold decoration | Caco-2 Papp A>B | 491 | 0.090 | 0.094 | 2.04 | 2.15 |
| Scaffold decoration | Caco-2 Efflux | 491 | 0.105 | 0.123 | 1.57 | 1.88 |
| Scaffold decoration | MPPB | 215 | 0.103 | 0.123 | 1.14 | 1.33 |
| Scaffold decoration | MBPB | 166 | 0.132 | 0.201 | 2.06 | 2.23 |
| Scaffold decoration | MGMB | 58 | 0.085 | 0.109 | 2.70 | 2.36 |

Note: the prediction coefficient of variation is reported as the median across groups because most stereoisomer groups receive near-identical predictions; the consistency ratio is reported as the mean because group-wise medians are dominated by zeros and hide the small non-zero tail that carries the stereochemical signal. Mean consistency ratios > 1 indicate amplification of within-group variation (scaffold-decoration groups for all endpoints except LogD under XGBoost, and all 9 endpoints under CheMeleon); values < 1 indicate smoothing (all stereoisomer endpoints except KSOL under XGBoost, and all 9 endpoints under CheMeleon).

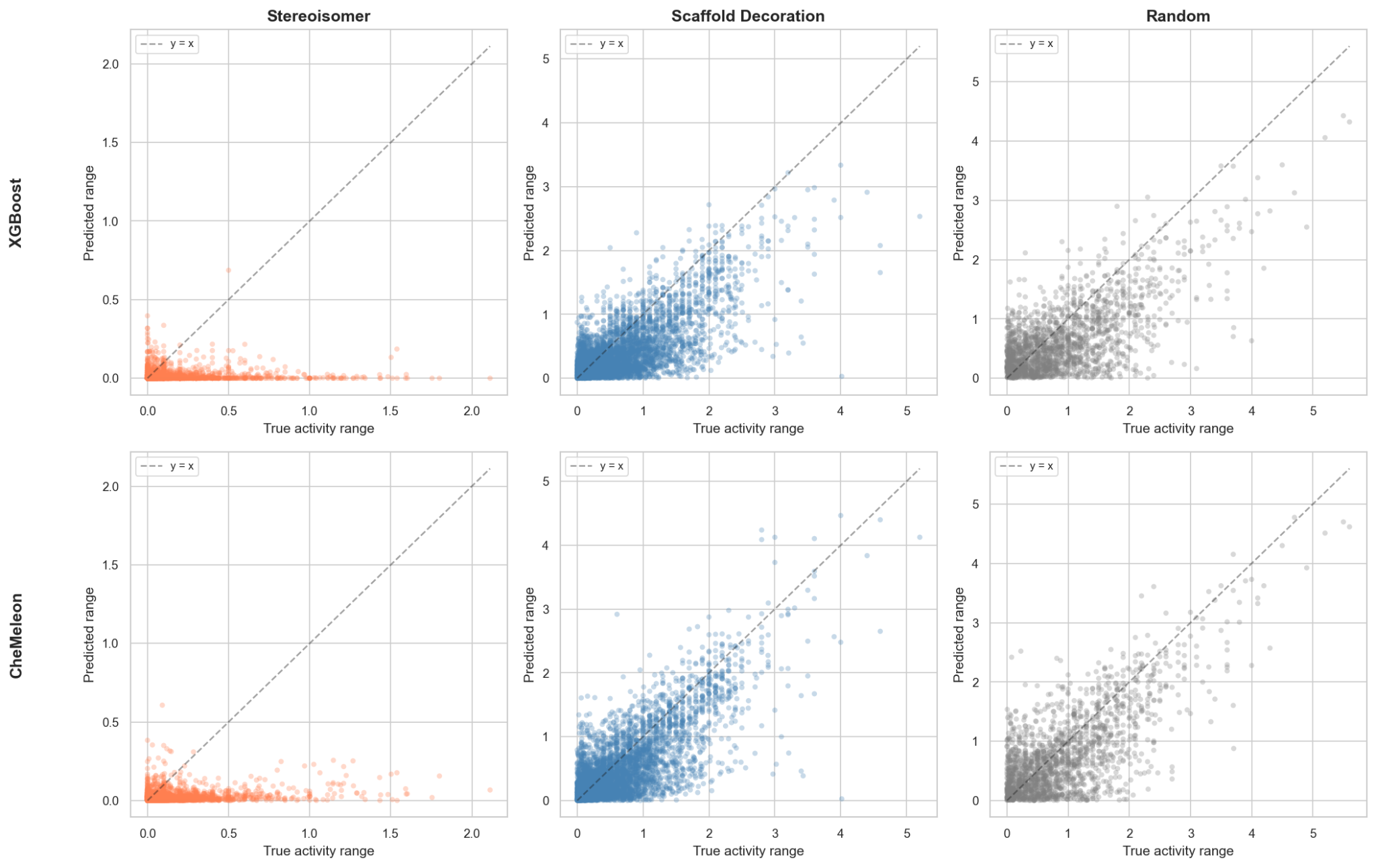

**Figure S11.** Within-group predicted range versus true activity range for each variant type, comparing XGBoost (top row) and CheMeleon (bottom row). Columns correspond to stereoisomer, scaffold decoration, and random groups. Points above the diagonal indicate that the model amplifies within-group differences; points below indicate smoothing. Stereoisomers cluster near the x-axis under both models, with small predicted ranges relative to true ranges, confirming that the stereoisomer under-prediction failure mode persists despite CheMeleon's graph-based stereo sensitivity. Scaffold decoration groups straddle the diagonal, and random pairs scatter broadly around it.

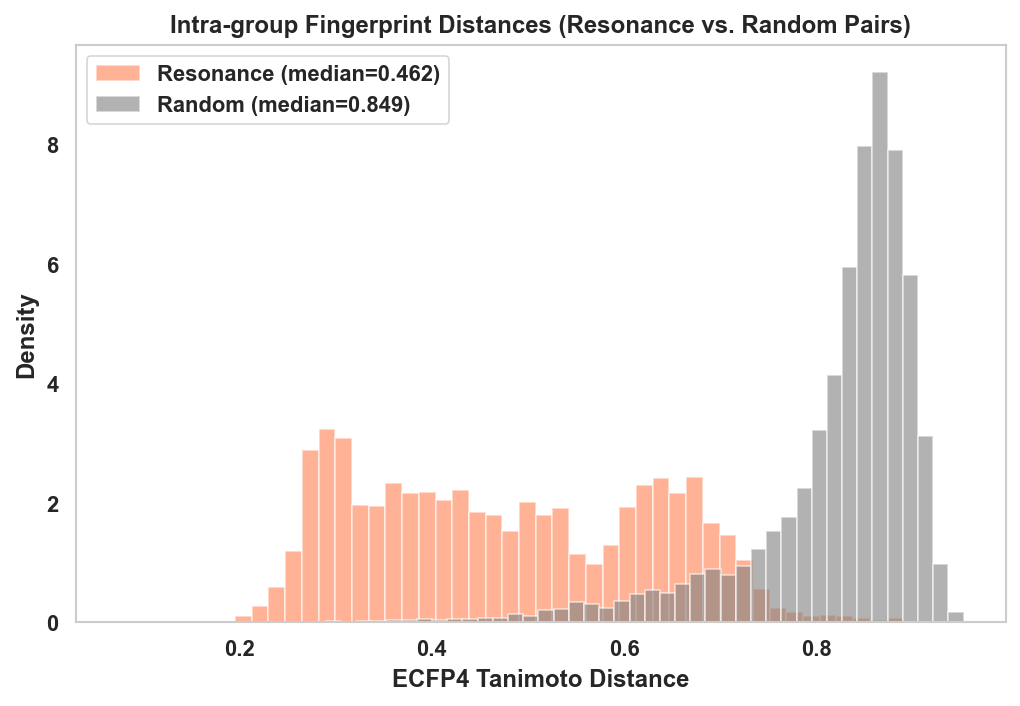

**Figure S12.** Intra-group Morgan fingerprint Tanimoto distance distributions for resonance forms (median 0.46) compared to random pairs (median 0.85). Resonance forms of the same molecule are far apart in fingerprint space despite being chemically identical.

**Table S6.** Resonance form RMSE impact by endpoint, ordered by XGBoost total swing, with side-by-side XGBoost and CheMeleon results. n_multi is the number of molecules with >1 distinct resonance form at the endpoint-relevant pH. Baseline RMSE uses the standard input representation; Improve % is the RMSE decrease from choosing each molecule’s lowest-error resonance form; Worsen % is the RMSE increase from choosing each molecule’s highest-error resonance form; Swing % is the total best-to-worst RMSE span relative to baseline.

| **Endpoint** | **n_multi** | **% affected** | **Baseline RMSE XGB** | **Baseline RMSE CheM** | **Improve % XGB** | **Improve % CheM** | **Worsen % XGB** | **Worsen % CheM** | **Swing % XGB** | **Swing % CheM** |
| --- | --- | --- | --- | --- | --- | --- | --- | --- | --- | --- |
| LogD | 2,060 | 28.2 | 0.60 | 0.47 | 7.0 | 4.3 | 20.4 | 16.3 | 27.4 | 20.6 |
| MLM CLint | 1,581 | 27.8 | 0.54 | 0.52 | 3.5 | 3.0 | 9.1 | 4.1 | 12.6 | 7.1 |
| MBPB | 367 | 25.7 | 0.26 | 0.24 | 3.7 | 2.1 | 7.6 | 9.8 | 11.3 | 11.9 |
| KSOL | 2,060 | 28.2 | 0.54 | 0.54 | 4.0 | 4.1 | 6.4 | 5.0 | 10.3 | 9.1 |
| HLM CLint | 888 | 19.6 | 0.51 | 0.51 | 1.7 | 1.8 | 7.3 | 2.9 | 9.0 | 4.7 |
| MPPB | 456 | 26.0 | 0.29 | 0.28 | 3.4 | 2.9 | 4.9 | 3.4 | 8.3 | 6.3 |
| MGMB | 230 | 53.4 | 0.27 | 0.27 | 1.8 | 1.3 | 3.8 | 4.3 | 5.7 | 5.7 |
| Caco-2 Papp A>B | 68 | 1.8 | 0.32 | 0.34 | 0.5 | 0.4 | 1.1 | 0.8 | 1.5 | 1.2 |
| Caco-2 Efflux | 68 | 1.8 | 0.32 | 0.33 | 0.2 | 0.3 | 0.2 | 0.3 | 0.4 | 0.5 |
